# Chronic social defeat stress induces meningeal neutrophilia via type I interferon signaling

**DOI:** 10.1101/2024.08.30.610447

**Authors:** Stacey L. Kigar, Mary-Ellen Lynall, Allison E. DePuyt, Robert Atkinson, Virginia H. Sun, Joshua D. Samuels, Nicole E. Eassa, Chelsie N. Poffenberger, Michael L. Lehmann, Samuel J. Listwak, Ferenc Livak, Abdel G. Elkahloun, Menna R. Clatworthy, Edward T. Bullmore, Miles Herkenham

## Abstract

Animal models of stress and stress-related disorders are also associated with blood neutrophilia. The mechanistic relevance of this to symptoms or behavior is unclear. We used cytometry, immunohistochemistry, whole tissue clearing, and single-cell sequencing to characterize the meningeal immune response to chronic social defeat (CSD) stress in mice. We find that chronic, but not acute, stress causes meningeal neutrophil accumulation, and CSD increases neutrophil trafficking in vascular channels emanating from skull bone marrow (BM). Transcriptional analysis suggested CSD increases type I interferon (IFN-I) signaling in meningeal neutrophils. Blocking this pathway via the IFN-I receptor (IFNAR) protected against the anhedonic and anxiogenic effects of CSD stress, potentially through reduced infiltration of IFNAR^+^ neutrophils into the meninges from skull BM. Our identification of IFN-I signaling as a putative mediator of meningeal neutrophil recruitment may facilitate development of new therapies for stress-related disorders.

**One sentence summary:** Type I interferon sensing neutrophils accumulate in meninges of psychosocially stressed mice via skull bone marrow channels and are associated with the negative behavioral sequelae of stress; blockade of this pathway inhibits neutrophil trafficking and improves behavioral outcomes.

## Introduction

Chronic inflammation has been linked to both psychosocial stress and major depressive disorder (MDD)^1,2^, highlighting a possible role for the immune system as an intermediary between psychological risk factors and the development of mood disorders. Proinflammatory cytokines produced by innate immune cells have attracted attention given robust data that they are associated with depression and depressive-like behavior in animal models^3,4^, including type I interferons (IFN-I), which induce depressive symptoms in otherwise nondepressed individuals^5–7^ and are associated with development of MDD^8,9^.

In parallel, multiple clinical studies have documented an increased ratio of neutrophils relative to lymphocytes in people with MDD^10,11^. Neutrophils are IFN-I-sensing phagocytes of the innate immune system that play a critical role as sentinels, acting as first responders to both pathogens and sterile injury^12–14^. Blood neutrophil levels rapidly elevate following acute stress via direct sympathetic innervation of bone marrow (BM)^15,16^, and stress is strongly associated with the development of depression^17^. We have furthermore demonstrated in MDD patients that peripheral neutrophil levels are the major immune cell subset most predictive of symptom severity^18^, suggesting a potential role for neutrophils in the pathogenesis of depression.

Mechanisms by which neutrophils contribute to depressive symptoms are currently unclear, however, we and others have demonstrated that cells acting from within the meninges play a key role in maintenance of behavioral homeostasis^19–21^. Specifically, we have shown that transgenic *Cd19^-/-^* mice, which are B cell deficient, are more anxious than wild type (WT) littermates at baseline^21^. Subjecting WT mice to chronic social defeat (CSD) stress–which reliably induces depressive- and anxiety-like behaviors^22–24^–led to decreased B cell numbers in the meninges. Bulk transcriptomic analysis of meningeal tissue from CSD-stressed and *Cd19^-/-^* mice revealed a shared increase in IFN-I signaling. Moreover, *Cd19^-/-^* mice showed elevated levels of meningeal neutrophils^21^.

Traditionally, neutrophils are thought to enter tissue from the bloodstream following BM egress. However, a brain-specific mechanism wherein neutrophils traffic directly via skull BM reservoirs to the meninges has recently been characterized^25^. These skull-derived neutrophils are preferentially recruited to the brain following tissue damage and in disease^26–28^ but their role in stress-related processes is unknown.

We hypothesized that neutrophils within the meningeal space were related to stress-induced depressive symptoms, and that these neutrophils originate from skull BM. We characterized immune cell dynamics and phenotypic changes in the meninges and in a variety of peripheral tissues following CSD stress. Using a data-driven approach, we identified CSD meningeal neutrophils as the source of enhanced IFN-I signaling previously seen in CSD^21^. This led us to hypothesize that IFN-I signaling promotes neutrophil egress from skull BM to the meninges, and in turn depressive- and anxious-like behaviors following CSD stress. Our preliminary evidence suggests that systemic IFN-I depletion normalizes the effects of CSD stress on meningeal neutrophils and behavior. This suggests that skull-to-meningeal trafficking of IFN-I sensing neutrophils is an important factor in the progression of CSD stress-induced behavioral changes.

## Results

### CSD increases meningeal and blood neutrophil abundance

Flow cytometry analyses on meningeal tissue from C57BL/6J WT mice showed that CSD increases meningeal neutrophil abundance (**Figures 1A**, **S1A-B**). Intravenous anti-CD45 antibody injections (CD45iv) were used to distinguish between parenchymal (iv^-^) and circulating blood-derived (iv^+^) cells (**Figures 1B, S1C-D**). There was a 1.3-fold increase in iv^-^ meningeal neutrophils (**Figure 1C**: \*\**p* < 0.01, *t* = 3.4, *df* = 47), but no increase in iv^+^ meningeal blood neutrophils (**Figure S1E**). There was a large, 5.3-fold increase in blood neutrophils (**Figure 1D**: \*\*\*\**p* < 0.0001, *U* = 36). We confirmed that the relative changes in neutrophil numbers (normalized to total CD45^+^ cells) were consistent with absolute count data in meninges (\*\**p* < 0.01, *t* = 3.2, *df* = 16) and blood (\**p* < 0.05, *t* = 3.0, *df* = 10). We also noted a CSD-related increase in meningeal monocytes (\*\**p* < 0.01, *U* = 132) and, as previously reported^21^, a decrease in meningeal B cells (\*\**p* < 0.01, *t* = 3.2, *df* = 47). See **Figure S2**.

**Figure 1:**
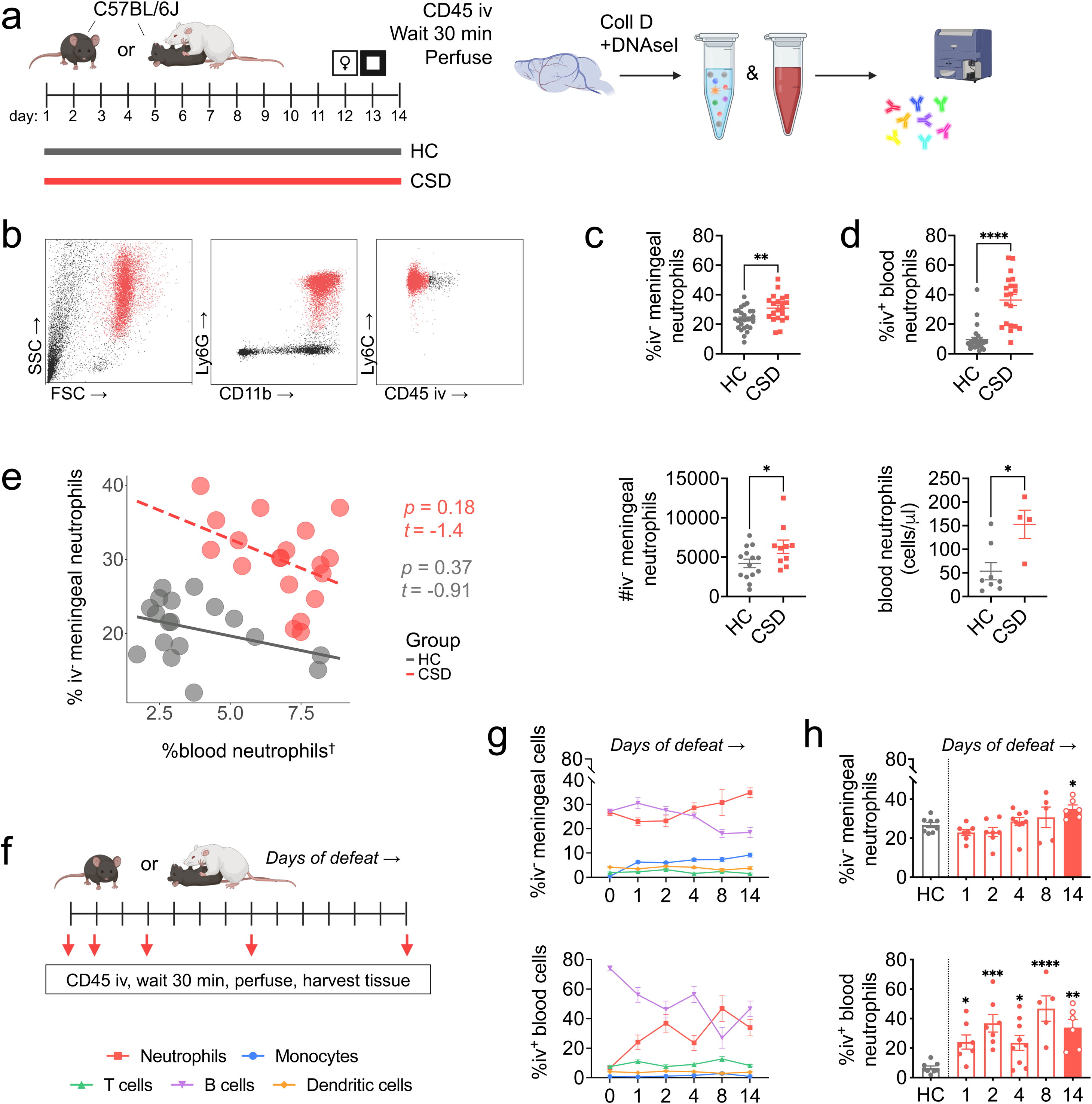
Meningeal neutrophils are elevated following chronic, but not acute, social defeat stress. **a**) CSD mice were behaviorally phenotyped on days 10-13, and tissue harvested at day 14 (**Figures S1A-C**). **b**) Mice were injected retro-orbitally with a fluorescently labeled CD45 antibody for exclusion of blood-exposed cells in the meninges. Simplified gating strategy showing identification of nonvascular neutrophils from meningeal tissue. Full gating strategy shown in **Figures S1D-E**. **c**) Flow cytometric analysis shows CSD stress causes an increase in meningeal neutrophils, defined as CD45iv^-^;CD11b^+^;Ly6G^+^;Ly6C^int^ and (*top)* calculated as a percent of live CD45^+^ cells (n_HC_ = 27, n_CSD_ = 21), or *(bottom)* assessed for absolute, as opposed to relative, cell counts (n_HC_ = 14, n_CSD_ = 10). **d**) Flow cytometry analysis of blood neutrophils (CD45iv^+^;CD11b^+^;Ly6G^+^;Ly6C^int^) shows a robust increase following CSD stress in both the (*top*) percentage of neutrophils (n_HC_ = 27, n_CSD_ = 20) and *(bottom)* absolute counts (n_HC_ = 8, n_CSD_ = 4). **e**) No apparent relationship between blood and meningeal neutrophils; ^†^values were square root transformed to improve normality. **f**) Schematic of acute vs chronic stress study; red arrows indicate time points in days at which mice were killed and tissue was harvested. **g**) Comparison of all cell types examined in this study showing equal and opposite directional fluctuations for neutrophils and B cells in both tissues. **h**) There was a main effect of the number of encounters for meningeal neutrophils, but only the CSD day-14 group showed a significant increase by post hoc analysis. In contrast, there was a significant increase in blood neutrophils overall and at each time point when compared to HC (subscript indicates days of defeat: n_HC_ = 8, n_1_ = 7; n_2_ = 7; n_4_ = 9; n_8_ = 5; n_14_ = 6). Diagrams made with Biorender. HC = home cage, CSD = chronic social defeat stress. Data shown as mean ± SEM. \**p* < 0.05, \*\**p* < 0.01, \*\*\**p* < 0.001, \*\*\*\**p* < 0.0001.

We tested the extent to which meningeal neutrophil numbers reflect peripheral blood neutrophil levels, modeling the effect of condition (HC vs. CSD) and blood neutrophils on meningeal counts (**Figure 1G**). We found no relationship between blood and iv^-^ meningeal neutrophils.

### Chronic but not acute stress induces meningeal neutrophil accumulation

In both blood and meningeal tissue, we observed dynamic changes in neutrophil and B cell populations with increasing exposure to social defeat (**Figures 1F-G, S2C**). While a single day of defeat stress was sufficient to elevate neutrophils in blood (\*\*\**P* < 0.001, *H* = 21.8; post hoc, \**p* < 0.05, *Z* = 2.3), no stress-induced elevation in meningeal neutrophils was observed until the day 14 time point (**Figure 1H**: \**P* < 0.05, *H* = 13.7; post hoc \**p* < 0.05, Z = 2.3). In contrast, there was a significant increase in blood neutrophils overall and at each time point when compared to HC (subscript indicates days of defeat: \**p_1_* < 0.05, Z_1_ = 2.3; \*\*\**p_2_* < 0.001, Z_2_ = 2.5; \**p_4_* < 0.05, Z_4_ = 2.3; \*\*\*\**p_8_* < 0.0001, Z_8_ = 4.0; \*\**p_14_* < 0.01, Z_14_ = 3.2). A decrease in meningeal B cells was apparent by the 8^th^ day of defeat (\*\*\**P* < 0.001, *H* = 22.3; post hoc \*\*\*\**p* < 0.0001, Z = 2.9), preceding the increase in meningeal neutrophils, though in blood the decrease in B cells was delayed (**Figure S2C**).

### Immunohistochemistry (IHC) replicates CSD-induced increase in meningeal neutrophils

Using LysM*^gfp/+^* mice, which strongly express GFP in neutrophils (**Figures 2A-B**), we replicated the effect of CSD stress on meningeal neutrophils. Confocal microscopy of meningeal whole mounts showed broad distribution of neutrophils both inside and outside blood vessels (**Figures 2C**-**D**). Quantitation revealed a 1.3x-fold increase in the number of meningeal neutrophils in CSD vs control (**Figure 2E**: \**p* < 0.05, *t* = 2.2, *df* = 23) - consistent with our flow cytometry data. The meningeal neutrophilia in CSD mice was observed across intravascular, abluminal (≤10μm away from a blood vessel) and parenchymal (>10μm away from a blood vessel) compartments (**Figure 2F**: \*\**p* < 0.01, *F*_(1,71)_ = 9.4; post hoc \**p* < 0.05, *q =* 4.3).

**Figure 2:**
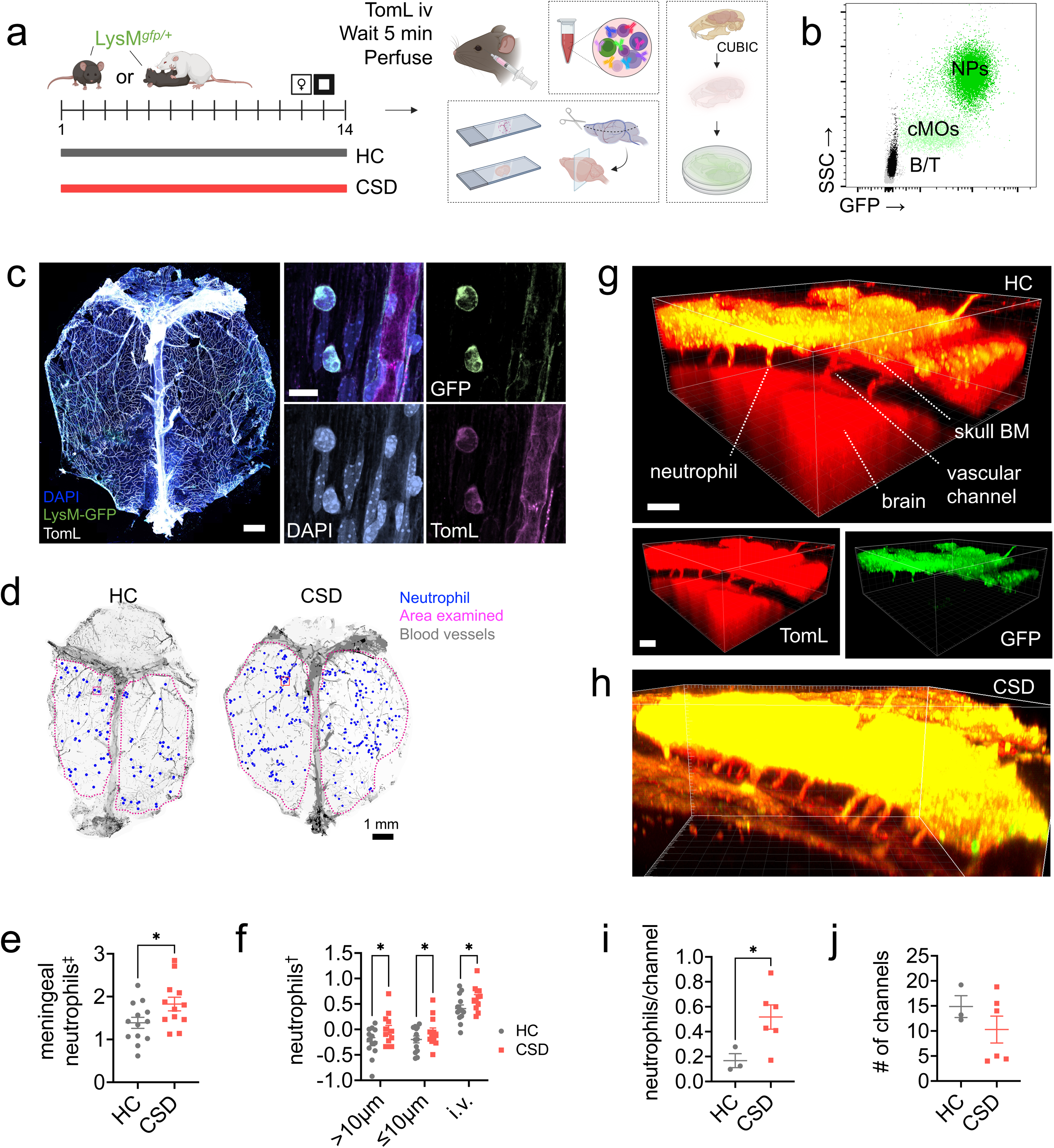
CSD stress leads to increased numbers of neutrophils in vascular channels connecting skull bone marrow to meninges. **a**) LysM*^gfp/+^* mice were subjected to CSD stress. Before TomL intravascular (iv) injection, blood was drawn for flow cytometry (**Figure SF3**). Dorsal meninges and brain (**Figure S4**) were prepared for imaging. A separate cohort of mice was used for tissue clearing. Diagram made with Biorender. **b**) Flow cytometry data of blood shows both high GFP expression and SSC in neutrophils compared to other cell types (x axis is log scale, y axis is linear). **c**) *Left*: Representative image for dorsal whole-mount meningeal tissue. Scale bar = 1 mm. *Right*: GFP^+^ neutrophils adjacent to a blood vessel; merge and individual channels. Scale bar = 10μm. **d**) Representative meningeal whole mounts from HC and CSD mice showing hand-counted meningeal neutrophils, normalized to indicated area. Blood vessels traced in Adobe Illustrator for visualization. Inset box represents approximate size of area shown in high resolution images from **(c)**. **e**) Quantification of total meningeal neutrophils shows an increase with CSD stress. ^‡^ natural log-transformed. **f)** Neutrophils were separated into three categories based on their location in the tissue: >10μm away from a blood vessel (“parenchymal”), ≤10μm away from a blood vessel (“abluminal”), and intravascular (“iv”). CSD led to neutrophil elevations in all three subcompartments. ^†^log_10_ -transformed. **g)** Representative image from cleared HC skull showing vascular channels between skull bone marrow and the meninges. Scale bar = 100μm. *Top*: Merged image. *Bottom*: Individual channels for blood vessels and neutrophils. **h)** Representative image from a CSD mouse. **i)** Quantification of neutrophils per channel (normalized to number of channels) indicates increased egress from skull bone marrow following CSD stress (n_HC_ = 3, n_CSD_ = 6). **j)** No differences in the number of channels counted between groups. HC = home cage, CSD = chronic social defeat stress, TomL = tomato lectin, NPs = neutrophils, cMOs = classical monocytes, SSC = side scatter complexity, BM = bone marrow. Data shown as mean ± SEM. \**p* < 0.05.

Otherwise, LysM*^gfp/+^* mice phenocopy C57BL/6J mice, recapitulating the expected effects of stress via a typical depressive/anxious behavioral response to the CSD stress paradigm, i.e., decreased urine scent marking (USM) preference (**Figure S3A**: \*\*\**p* < 0.001) and decreased open field (OF) crosses to center (**Figure S3B**: \*\**p* < 0.01, *t* = 3.27, *df* = 2), respectively. CSD also induced blood neutrophilia (**Figure S3C**: \*\*\**p* < 0.001, *U* = 12), blood monocytosis (**Figure S3D**: \*\**p* < 0.01, *U* = 28.5), and blood lymphopenia – i.e., decreases in blood T cells (**Figure S3E**: \**p* < 0.05, *U* = 21), and blood B cells (**Figure S3F**: \*\**p* < 0.01, *U* = 18).

IHC examination of brain tissue revealed no evidence for neutrophil infiltration into brain parenchymal tissue, though there were significantly more neutrophils ‘stuck’ in the neurovasculature in CSD mice (**Figures S4A-B**: \*\*\*\**p* < 0.0001, *F*_(1,52)_ = 22.1. post hoc, *q* = 6.7, \*\*\**p* < 0.001 in medial prefrontal cortex, striatum, and hippocampal sections).

### CSD increases neutrophil trafficking between skull BM and meninges

We used tissue clearing to visualize GFP^+^ cells in the channels connecting skull BM to the meninges (**Figures 2G-H**). There were more GFP^+^ neutrophils per vascular channel in CSD mice compared to HC (**Figure 2I**: \**p* < 0.05, *t* = 2.4, *df* = 7), suggesting this route may be important for neutrophil trafficking into the meninges.

### Increased neutrophil levels correlate with stress-related behavioral changes

Increased iv^-^ neutrophils from C57BL/6J meningeal tissue were associated with increasing USM-assayed anhedonia (**Figure 3**: *FDR < 0.05, z = −2.4, **Table S1**). Parenchymal neutrophils from LysM*^gfp/+^* mice showed a similar relationship and effect size on USM behavior (**Figure S5**: * unadjusted *P* < 0.05, z = −2.0, **Table S3**). In the LysM*^gfp/+^* cohort, there was also an association between increasing anxiety-like behavior in the OF task and increased parenchymal (*FDR < 0.05, t = −3.1) and abluminal (*FDR < 0.05, t = −2.6) neutrophil levels (**Figure S5**, **Table S4**). A similar trend with iv^-^ neutrophils was observed in C57BL/6J mice (**Figure 3**; *P* = 0.17, t = −1.43, **Table S2**).

**Figure 3:**
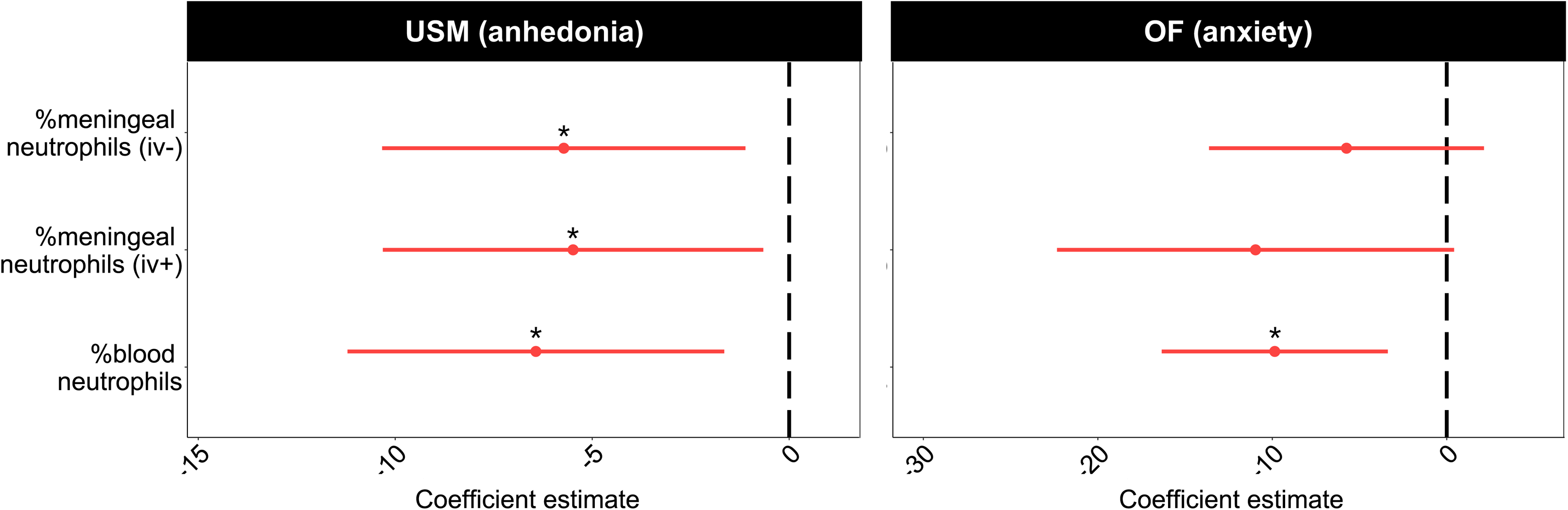
Elevated neutrophil levels are associated with the negative behavioral sequelae of CSD stress. Dot and whiskers plots showing standardized effect sizes for neutrophil levels on behavioral outcomes (batch corrected for cohort). *Left*: Logistic regression models show negative relationships between blood and meningeal neutrophils with USM behavior. As neutrophil levels rise, mice are less likely to engage with the hedonic stimulus, i.e., odor from sexually mature females (n_HC_ = 12, n_CSD_ = 13). *Right*: Linear regression models indicate blood neutrophils are significantly associated with anxiety-like behavior. Specifically, as blood neutrophil levels increase, mice explored the OF arena less (*iv^-^/iv^+^ meningeal neutrophils:* n_HC_ = 15, n_CSD_ = 17; *blood:* n_HC_ = 10, n_CSD_ = 11). See **Tables S1**-**S2** for full statistics; see **Figure S5** for comparison with LysM*^gfp/+^* mice. HC = home cage, CSD = chronic social defeat stress, USM = urine scent marking, OF = open field. \**p* < 0.05.

Peripherally, increased blood neutrophil levels showed a significant relationship with both USM-assayed anhedonic behavior (*FDR < 0.05, z = −2.6) and anxiety-like behavior on OF (*FDR < 0.05, t = −2.98); see **Figure 3**, **Tables S1-S2**. Similar patterns were seen in LysM*^gfp/+^* mice (**Figure S5, Tables S5-S6**) – blood neutrophils trended toward an association with both anhedonia (unadjusted *P* = 0.11, z = −2.6) and anxiety-like behavior (unadjusted *P* = 0.065, t = −1.99). Increased circulating B cells were associated with decreased anxiety-like behavior (**Figure S5**: * unadjusted *P* < 0.05, t = 2.2, **Table S6**), consistent with our previously published data^21^.

### Single cell RNA sequencing (scRNAseq) reveals increased neutrophils in CSD meninges, increased proinflammatory signaling, neutrophil heterogeneity

scRNAseq was used to profile meningeal immune cells from behaviorally stratified HC and CSD mice (**Figures 4A-B**). Clustering (leidenalg; see **Supplementary Methods**) identified 20 cell clusters which were manually annotated based on expression of canonical lineage genes (**Figures 4C**, **S6A**-**C**). There was a large cluster of neutrophils, identified by high *Cxcr2* expression, which did not appear to be undergoing local cell proliferation (**Figures S6D**-**E**). Consistently with our IHC and flow cytometric data, the meninges from CSD-stressed mice showed a relatively greater proportion of neutrophils compared to HC (**Figure 4C**, lavender bars).

**Figure 4:**
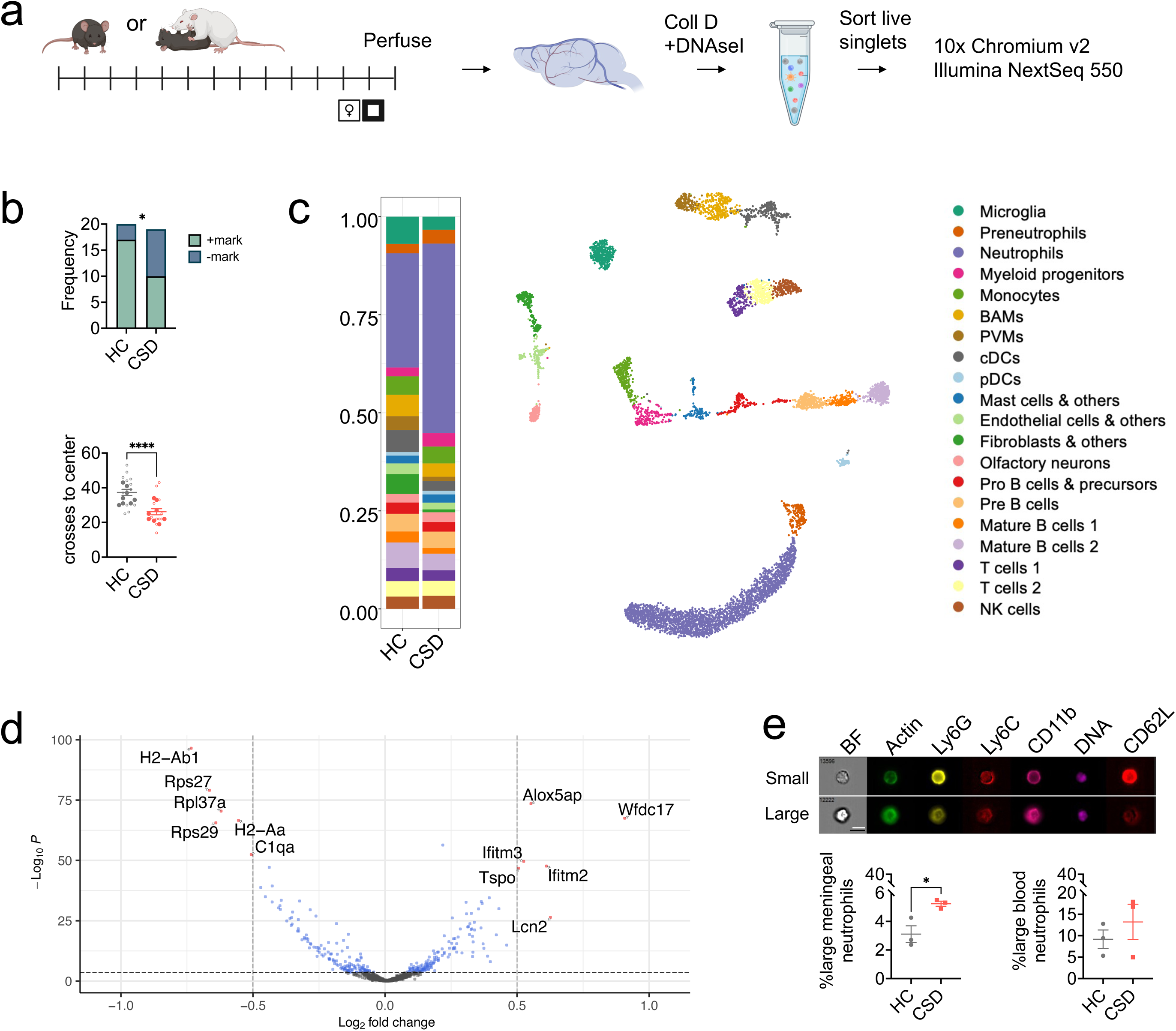
Single cell sequencing of meningeal tissue validates CSD neutrophilia; differential gene expression (DGE) analysis reveals enrichment of pathways related to cytoskeletal processes. **a**) Timeline for sample collection and processing. Figure made with Biorender. **b**) WT mice were behaviorally phenotyped prior to single cell analysis; a subset with representative behavior in both the USM task for anhedonia and the OF task for anxiety-like behavior were selected. *Top*: HC animals that marked (+mark) and CSD animals that did not mark (-mark) were chosen. *Bottom*: larger, filled circles indicate individual animals used for single-cell analysis. **c**) Analysis of 10x Genomics single-cell data for meningeal tissue reveals 20 distinct immune cell clusters. *Left:* Visualization of recovered cells as a proportion of total cells recovered per group; lavender indicates neutrophils. (n_HC_ = 8, n_CSD_ = 4; see **Methods**). Plot shown previously^21^. **d**) Volcano plot showing DGE between CSD and HC in the neutrophil cluster (excluding preneutrophil cluster). Indicated points represent DGE with LFC > 0.5 and FDR *p* < 0.001. Gene set enrichment analysis (GSEA) revealed several enriched pathways related to cell size and the cytoskeleton (see **Figure S8**). **e)** The Amnis ImageStream system was used to visualize cells stained with flow cytometry markers identifying neutrophils, phalloidin (to label actin) and Hoechst (to label nuclei). *Top*: Representative images acquired from individual meningeal neutrophils. Two differently sized populations emerged, as depicted (white bar = ‘small’ neutrophil diameter, black bar = ‘large’). *Bottom*: There were nearly 3x more enlarged neutrophils in CSD meninges compared to HC. No changes were evident in blood. For more details, see **Figure S9**. HC = home cage, CSD = chronic social defeat stress, BAM = border associated macrophage, PVM = perivascular macrophage, DBC = dural border cell, BF = brightfield. Data shown as mean ± SEM. \**p* < 0.05, \*\*\*\**p* < 0.0001.

Subclustering of neutrophils alone identified 6 subclusters with roughly equal proportions of each subtype in both HC and CSD meninges (**Figure S7A**). Based on expression of genes related to primary, secondary, and tertiary granule formation, as well as suppression of transcriptional machinery (**Figure S7B**), these subclusters likely correspond to states of increasing neutrophil maturation^29–31^. Pseudotime analysis of differentially expressed genes across the subclusters supported this hypothesis (**Figure S7C**).

### Effects of CSD stress on neutrophil cell size and actin polymerization

Gene set enrichment analysis (GSEA) of differentially expressed genes (DEGs) in the pooled neutrophil cluster uncovered 80 biological pathways associated with CSD stress (**Figure S8**). 8 of the 41 positively enriched pathways indicated CSD was associated with altered neutrophil cell size and/or actin polymerization, e.g. the gene ontology (GO) pathway “regulation of cellular component size” (**Figures S9A-B**). Chronically high demand for neutrophils leads to the release from BM of immature cells^32,33^, which are both volumetrically larger^34^ and more rigid^35^ than mature neutrophils. Given this and our GSEA results, we hypothesized there would be an increase in enlarged BM-derived neutrophils in the meninges.

To test this, we analyzed neutrophils on an Amnis ImageStream imaging flow cytometer (**Figures 4E**, **S9C**). We verified the expected CSD increase in meningeal (\*\**p* < 0.01, *t* = 7.4, *df* = 4) and blood (\**p* < 0.05, *t* = 2.9, *df* = 4) neutrophils (**Figure S9D**), and identified a subset of relatively enlarged neutrophils in both tissues. There was a significant, ∼3x increase in the enlarged neutrophil subset in CSD meninges (\**p* < 0.05, *t* = 3.5, *df* = 4), but not in blood (**Figure 4E**). There were no group differences in actin MFI for meningeal neutrophils, but CSD led to increased actin MFI in circulating neutrophils (**Figure S9E**: \**p* < 0.05, *F*_(1,9)_ = 9.6).

Pathway analysis also implicated increases in the GO pathway “cytokine production” in stressed animals, with increased expression of *Cxcr2* in CSD neutrophils (**Figures S10A-B**). However, we did not detect changes in either the frequency (%CXCR2^+^) or protein expression (CXCR2 MFI) in any examined tissue (**Figures S10C-D**). We did observe an increased percentage of neutrophils in CSD meninges, skull, tibia, and blood (\*\*\*\**p* < 0.0001, *F*_(4,78)_ = 44.8; post hoc: *t* = 7.9, \*\*\*\**p* < 0.0001), consistent with our other findings and with stress-associated increases in BM neutrophil levels described elsewhere^15^. We also noted an increase in CSD CXCL1 plasma concentration (\*\**p* < 0.01, *U* = 9), apparently due to increased mRNA expression in the liver (\**p* < 0.05, *U* = 8), and decreased *Cxcl12* expression in CSD tibia (\**p* < 0.05, *t* = 2.3, *df* = 17). Collectively, this would promote egress of neutrophils into the blood stream, i.e. through CXCL1 binding to CXCR2 and decreased CXCL12 binding to CXCR4 (**Figure S10E**). Finally, detection of ROS by intracellular flow cytometry provided no evidence for increased neutrophil ROS production on a per cell basis, consistent with the GO pathway “cell redox homeostasis” (**Figure S11**).

### Enriched IFN-I signaling in meningeal neutrophils

Our GSEA results also identified enrichment of the GO pathway “response to type I interferons” (**Figures S8, S12A**), which we pursued further given extensive literature supporting a role for IFN-I in promoting depressive-like behavior^5,6,36^. Across all meningeal cell clusters, leading-edge genes *Interferon induced transmembrane proteins (Ifitm)-2* and *Ifitm3* showed highest levels of expression in neutrophils and monocytes (**Figure S12B**). *Ifitm2* and *Ifitm3* expression was strongly increased in CSD-stressed meningeal neutrophils generally (**Figures 4D**, **S12C**) and in several neutrophil subclusters (**Figures 5A-B**). In contrast, monocyte expression of *Ifitm2* and *Ifitm3* did not show stress-associated changes (**Figure S12C**).

**Figure 5:**
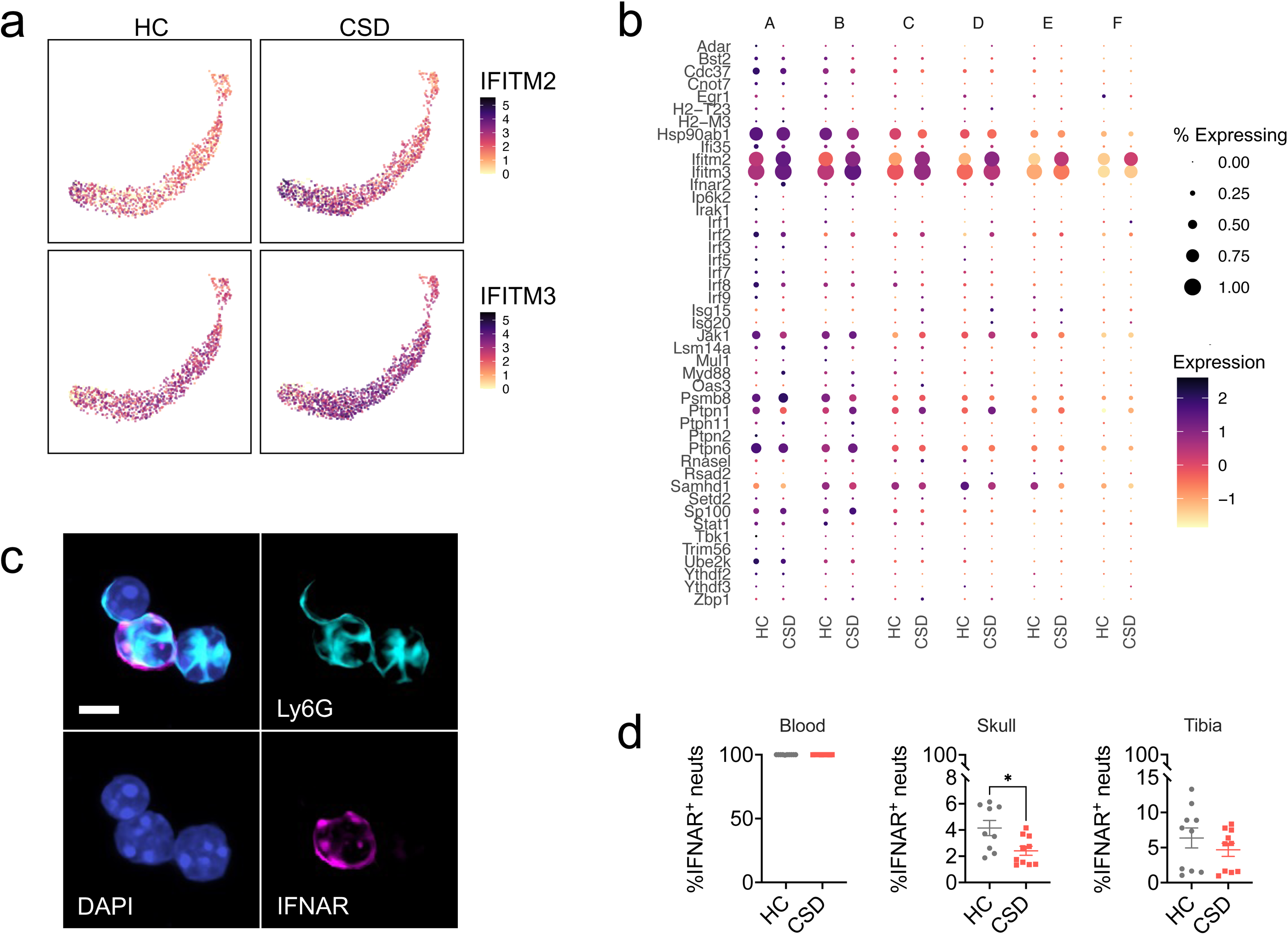
Migration of IFNAR^+^ neutrophils from skull bone marrow to the meninges may underlie the type I interferon neutrophil signature seen in CSD stressed mice. **a**) UMAP shows expression of *Ifitm2* and *Ifitm3*, the leading-edge genes for enrichment of the GO: “Response to type I interferon” pathway in neutrophils, in CSD compared to HC animals. **b**) Dot plot showing gene expression in each neutrophil subcluster for all genes comprising this pathway. Expression is scaled to mean ± standard deviation. **c**) 60x magnification of skull bone marrow-derived neutrophils. *Clockwise from top left*: Merge, Ly6G^+^ staining (cyan), IFNAR^+^ staining (magenta), nuclei (DAPI, blue). Scale = 5μm. **d**) IFNAR^+^ neutrophils, normalized to total neutrophils, for blood, skull, and tibia bone marrow. The population of IFNAR^+^ skull bone marrow neutrophils was decreased in CSD stressed mice, and may represent a migration event to the meninges. See **Figure S13** for more detail. HC = home cage, CSD = chronic social defeat. Data shown as mean ± SEM. \**p* < 0.05.

### IFNAR^+^ neutrophil egress from skull BM

IFN-I sensing meningeal neutrophils could migrate to meninges from peripheral BM tissue (via the circulation), or directly from skull BM^26–28^. We would predict a decrease in IFNAR^+^ neutrophils in the source tissue as they leave to traffic to the meninges. IFNAR^+^ neutrophils were present in blood, skull and tibial BM (**Figures 5C, S13A**). Comparison of HC vs CSD tissue revealed a 1.7-fold decrease of IFNAR^+^ neutrophils in CSD mice that was specific to skull BM (**Figure 5D**: \**p* < 0.05, *t* = 2.7, *df* = 17). This result, along with the neutrophils seen in skull-to-meninges vascular channels (**Figure 2I**), suggest skull BM as the source of stress-induced meningeal neutrophils.

### IFNAR^+^ neutrophils are a relatively mature pool of BM neutrophils

We noted three distinct populations of neutrophils – IFNAR^neg^, IFNAR^lo^, and IFNAR^hi^ – with tissue-specific distributions and expression of activation markers (**Figure S13**). In blood, the proportion of IFNAR^lo^ and IFNAR^hi^ neutrophils was approximately equal, with no detectable IFNAR^neg^ neutrophils. Bone marrow IFNAR^+^ neutrophils had higher side scatter complexity (SSC: skull, \*\*\*\**p* < 0.0001, *F*_(2,54)_ = 17.0; tibia, \*\*\*\**p* < 0.0001, *F*_(2,54)_ = 12.8)—a phenomenon corresponding to increased granule content and/or more complex nuclear morphology reflecting cell maturation. Conversely, the majority (>94%) of BM neutrophils were IFNAR^neg^, consistent with an expected reservoir of immature cells in this niche. We also examined MFIs in each of the three IFNAR neutrophil subtypes. Ly6C is expressed in response to IFN-I signaling^37^. Consistently, Ly6C MFI was highest in IFNAR^hi^ neutrophils from all three tissues (**Figure S13C**: blood, \*\*\*\**p* < 0.0001, *F*_(1,36)_ = 19.8; skull, \*\*\*\**p* < 0.0001, *F*_(2,54)_ = 19.7; tibia, \*\*\*\**p* < 0.0001, *F*_(2,54)_ = 16.2).

### IFNAR blockade improves the behavioral response to CSD stress

We assessed whether blockade of IFN-I signaling rescues the negative behavioral sequelae associated with CSD stress in LysM*^gfp^*^/+^ mice by administering an IFNAR-blocking antibody, as shown in **Figure 6A**. As expected, in the USM test for anhedonia, more HC+control antibody (IgG) mice marked compared to the CSD+IgG group (**Figure 6B**: \*\**p* < 0.01, χ^2^ = 9.5; post hoc \**p* < 0.05). Anti-IFNAR treatment rescued the phenotype, with no difference in marking frequency between HC+IgG and CSD+IFNAR groups. Anxiety data for the CSD+IFNAR blockade group was non-normally distributed, with distinct groups of animals that marked (+) or did not mark (-) in the USM test, so we treated these animals separately in our remaining analyses. There was a significant difference in the OF task between HC+IgG and CSD+IFNAR(-), where CSD+IFNAR(-) animals were more anxious. As before, behavior was rescued in the CSD+IFNAR blockade group, with no significant difference in behavior from HC+IgG (**Figure 6C**: \**P* < 0.05, *H* = 9.1; post hoc: \**p* < 0.05, *Z* = 2.8).

**Figure 6:**
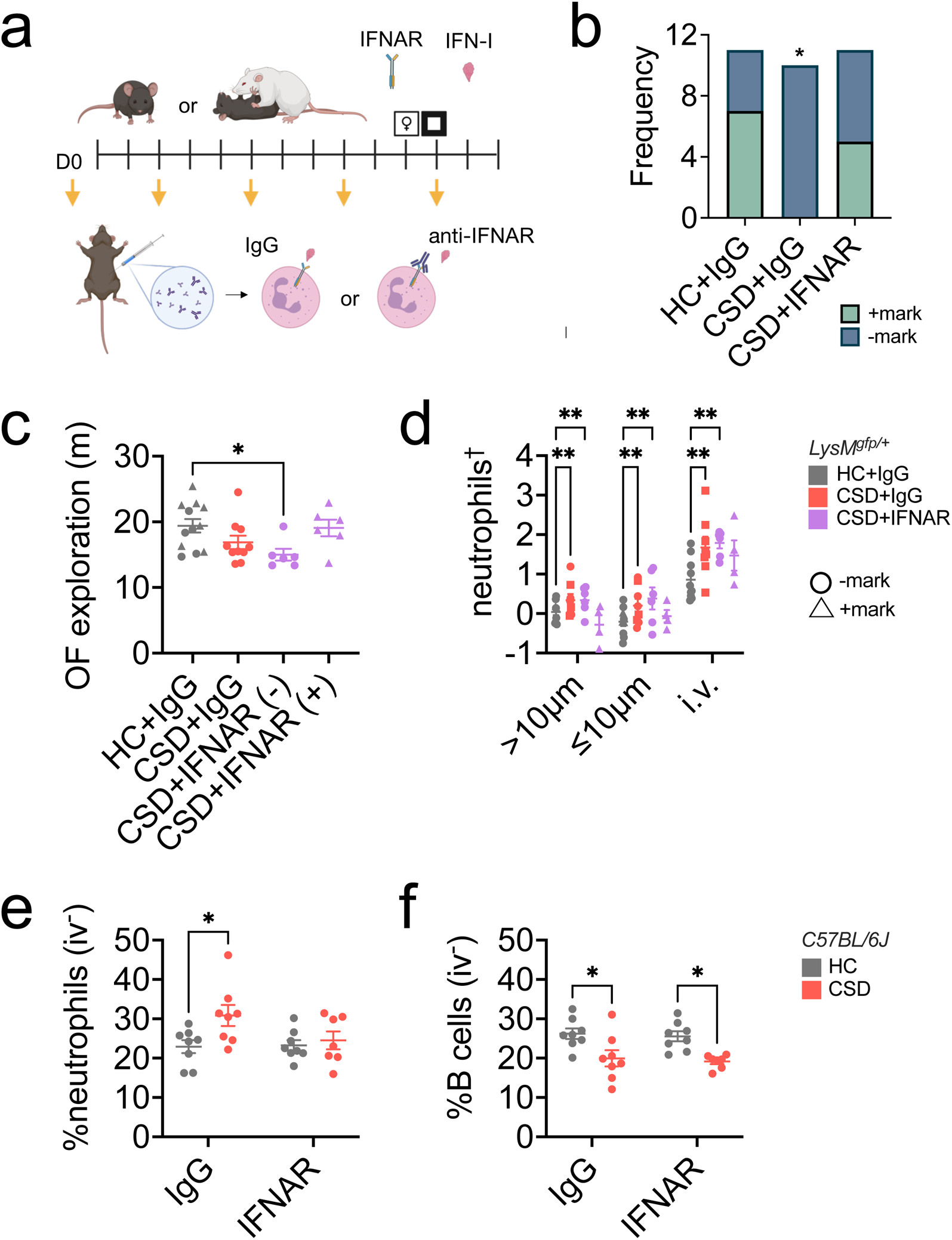
Type I interferon receptor (IFNAR) blockade may improve CSD-stress related behavioral anhedonia and prevent meningeal neutrophil accumulation. **a**) Schematic showing drug-delivery schedule for IFNAR blocking antibodies. Mice were injected with either matched IgG control antibody or anti-IFNAR antibody on the days indicated with yellow arrows. Figure made with Biorender. **b**) USM results from LysM*^gfp/+^* mice; as expected, CSD+IgG mice marked less frequently compared to the HC+IgG group. There were no differences between the HC+IgG group and the CSD+IFNAR group (n_HC+IgG_ = 11, n_CSD+IgG_ = 10, n_CSD+IFNAR_ = 11). Hereafter we considered LysM*^gfp/+^* mice in two separate groups – those that marked in the USM test (+, indicated with triangles) and those that didn’t (-, indicated with circles). **c**) HC+IgG(+) and (-) mice showed a normal distribution for anxiety-like behavior and were collapsed into one group. CSD+IFNAR(+) mice showed control-like levels of exploration in the OF task for anxiety-like behavior. **d**) There was an overall effect of group on meningeal neutrophils. Post hoc analysis indicated a CSD stress-induced increase that was normalized in the CSD+IFNAR(+) subgroup. ^†^values were natural log-transformed to improve normality. **e**) In C57BL/6J wild type mice there was an IFNAR-mediated rescue in meningeal neutrophil accumulation following CSD stress, with expected differences in IgG control groups (n_CSD+IFNAR_ = 7, otherwise n = 8). **f**) Anti-IFNAR treatment does not rescue the reduction in meningeal B cells seen with CSD stress. HC = home cage, CSD = chronic social defeat stress. Data shown as mean ± SEM. \**p* < 0.05, \*\**p* < 0.01.

### IFNAR blockade attenuates neutrophil migration into, but not B cell egress from, meningeal tissue in CSD stressed mice

Hand counting of meningeal neutrophils from these mice showed an expected increase in the CSD+IgG group, which was present irrespective of distance from blood vessel (**Figure 6D**: \*\*\**p* < 0.001, *F*_(3,78)_ = 7.9; post hoc \*\**p* < 0.01, *t* = 3.8). The CSD+IFNAR(-) group also showed an increase in meningeal neutrophils independent of distance from a blood vessel (\*\**p* < 0.01, *t* = 4.1). There was no significant difference between the HC+IgG group and the CSD+IFNAR(+) group; rather, there appeared to be a decrease in meningeal neutrophils >10 μm from a blood vessel in CSD+IFNAR(+) mice.

In WT mice, IFNAR blockade successfully attenuated neutrophil accumulation in the meninges (**Figure 6E**: \**p* < 0.05, *F*_(1,27)_ = 5.0) with the expected increase in control IgG antibody-treated CSD mice (\**p* < 0.05, *q =* 4.0). The rescue appeared to be specific to neutrophils (**Figure S14A**), though meningeal B cell depletion occurred in both antibody treatment groups for CSD stress (**Figure 6F**: \*\*\**p* < 0.001, *F*_(1,27)_ = 18.7).

Unexpectedly, there were no effects of anti-IFNAR treatment or CSD on peripherally circulating WBC populations, including both neutrophils and B cells (**Figure S14B**). Further examination of our data suggested an effect of repeated injection stress on circulating WBCs in non-defeated control mice (**Figure S15**). Importantly, no effect of injection stress was evident in the meninges.

### Reduced expression of chemorepellent factors in CSD stress may permit neutrophil entry into the leptomeninges

Expression of class 3 semaphorins was examined in microarray data from meningeal tissue. *Sema3b* expression was significantly reduced in CSD mice compared to HC (**Figure 7C**: LFC = −0.30, *unadjusted *P* = 0.050).

**Figure 7:**
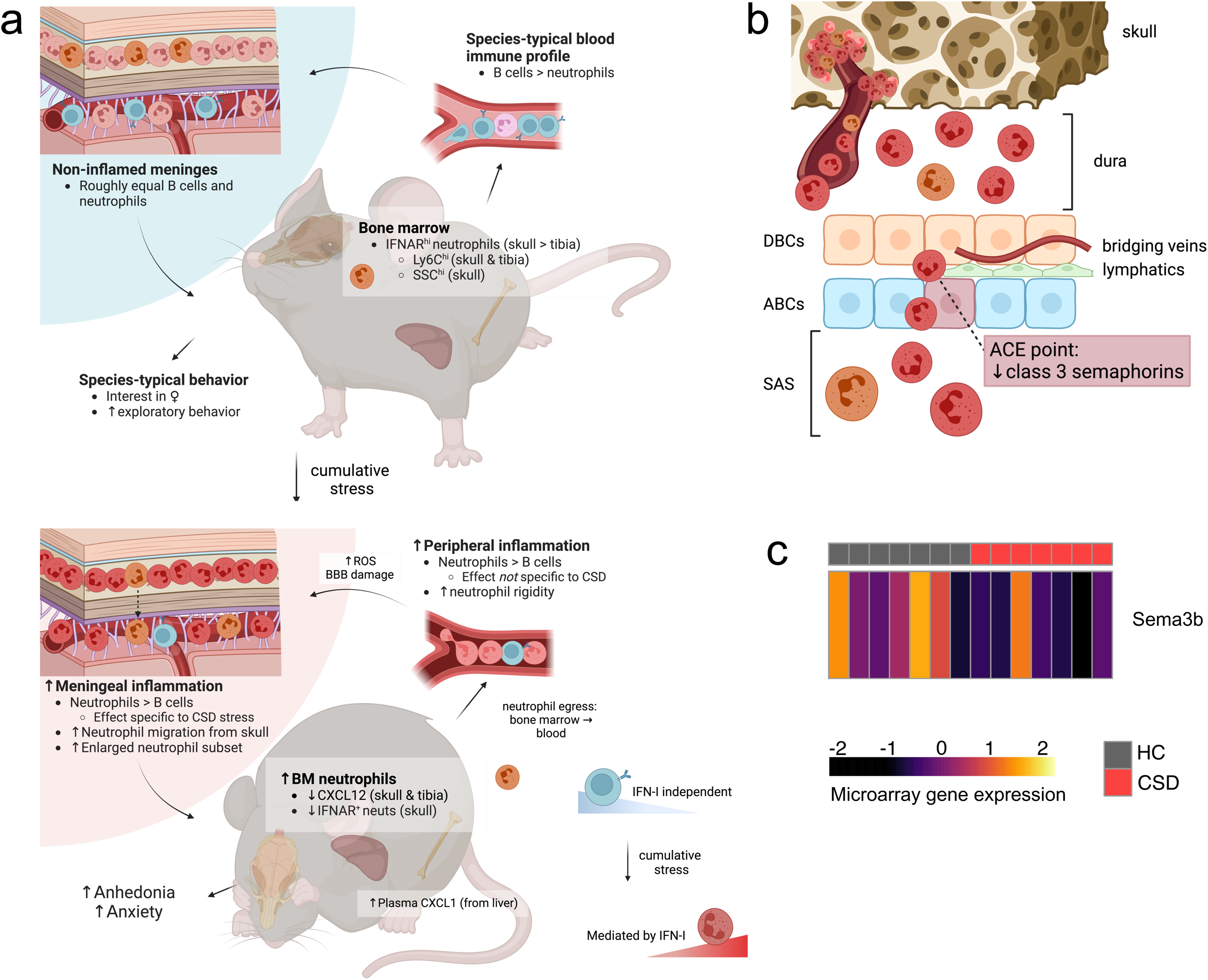
Proposed model for how chronic, but not acute, stress leads to dysregulation of the meningeal environment. **a**) At baseline, both skull and tibia bone marrow contain a small population of IFNAR^+^ neutrophils (orange), further divisible into IFNAR^lo^ and IFNAR^hi^ populations. Relatively mature IFNAR^hi^ cells express more Ly6C and may thus be more proinflammatory^37^. In blood, B cell numbers greatly outnumber neutrophils; in the meninges these two populations are roughly equal. Acute stress exposure leads to repetitive release of neutrophils into the blood but is not sufficient for accumulation of neutrophils in the meninges (Figures 1H, **S15A**). Prolonged exposure to psychosocial stress leads to a decline in meningeal B cells (**Figure S2C**) that precedes an increase in meningeal neutrophils (Figure 1H). Whereas CSD-associated meningeal neutrophilia appears to be mediated by IFN-I signaling, meningeal B cell depletion does not (Figures 6C,**E**). CSD stress causes random neurovascular damage^22^—potentially via stalled and rigid ROS-producing neutrophils in brain capillaries (**Figures S4, S9E, S11**)—which amplifies neutrophil recruitment from adjacent skull bone marrow (Figure 2G). **b**) Class 3 semaphorins (SEMA3) prevent migration of neutrophils across dural lymphatics into the leptomeninges at ACE points due to their chemorepellent properties^39^. Depletion of SEMA3 family expression leads to increased accumulation of neutrophils and other immune factors in SAS from whence they can directly interact with the brain to influence mood and behavior. **c**) Microarray analysis of SEMA3 family members in bulk meningeal tissue suggests decreased expression of *Sema3b*, which may permit neutrophil entry into the SAS (n = 7 per group). Figures made in Biorender. HC = home cage, CSD = chronic social defeat, SSC = side scatter complexity, ROS = reactive oxygen species, BBB = blood brain barrier, DBC = dural border cell, ABC = arachnoid barrier cell, ACE = arachnoid cuff exit, SAS = subarachnoid space.

## Discussion

Our data robustly demonstrate increased neutrophils in the meninges following CSD stress, and that neutrophilia correlates with a negative behavioral response to chronic stress (**Figures 3, S5**). Moreover, we have replicated the blood neutrophilia described in other preclinical stress models^15,38^ and in people with MDD^18^—though, interestingly, neutrophil levels in the meninges and blood were not significantly related (**Figure 1E**). These results are summarized in **Figure 7A**. Whereas acute stress led to a rapid increase in blood neutrophils (i.e., within 2 h of a single defeat encounter), the effect in meninges was quite specific—neither acute nor non-psychological stress recapitulated the phenotype (**Figures 1H, S15A**). This is consistent with a growing body of literature suggesting that the meningeal niche is governed by processes distinct from the peripheral immune system^21,25^.

It is not presently clear exactly how meningeal neutrophils influence behavior given no evidence for their entry into the brain parenchyma (**Figure S4**). However a mechanism for neutrophil entry into the subarachnoid space (SAS) via arachnoid cuff exit (ACE) points was recently described^39^; this occurs via reduced expression of chemorepellent class 3 semaphorins by arachnoid barrier cells (ABCs; **Figure 7B**). Any soluble factors released from neutrophils within this space have the capacity to interact directly with the brain. We reexamined our data from a previously published gene expression microarray on meningeal tissue^21^ in an attempt to address whether CSD stress influences semaphorin expression. *Sema3b* expression was reduced in CSD mice compared to HC (**Figure 7C**), suggesting permissive conditions for neutrophil entry into the SAS. Future efforts to demonstrate this pathway conclusively will be important.

We have previously shown that B cell depletion increases both IFN-I signaling and meningeal neutrophil levels in *Cd19^-/-^* mice^21^, which are deficient in B cells^40^. Here, we replicated the depleting effect of CSD stress on meningeal B cells and demonstrated that CSD stress leads to an enriched IFN-I signature in neutrophils but not in other meningeal cell types (**Figures 4E**, **S12**). Our unbiased identification of this pathway in multiple CSD cohorts using multiple experimental techniques is further strengthened by consistent evidence from humans, non-human primates, and rodents that IFN-I is sufficient to induce depression and depressive-like behavior^5,6,36^.

Differential gene expression (DEG) analysis of our single-cell RNA sequencing data indicated altered innate immune system activation across the neutrophil cluster (**Figure 4D**). For example, the most strongly upregulated transcript in CSD meningeal neutrophils was *Wfdc17*, which inhibits NFκB-mediated microglial activation^41^ and is proinflammatory in monocytes^42^. There was also evidence for altered proinflammatory leukotriene signaling: *Arachidonate 5-lipoxygenase-activating protein* (*Alox5ap*) is an accessory protein required for leukotriene synthesis, and *Translocator protein* (*Tspo*) mediates neutrophil chemotaxis via leukotriene B4 (*Ltb4*) receptor signaling^43,44^. Conversely, decreased *C1qa* expression following stress may hamper phagocytosis and clearance of dying neutrophils by nearby macrophages, leading to sustained local inflammation^45,46^.

Expression of *Ifitm2* and *Ifitm3,* the two leading-edge genes identified in CSD neutrophils from the “response to type I interferons” GO pathway, has been reported in a mature, hyperinflammatory neutrophil subtype observed in blood from humans and mice^30,47,48^. Characterization of IFN-I receptor (IFNAR) expressing neutrophils in different tissue compartments revealed highest Ly6C expression in IFNAR^hi^ neutrophils, which—in BM—appeared to also be more mature (**Figure S13C**). Consistently, qualitative assessment of IFNAR^+^ compared to IFNAR^-^ skull neutrophils showed more hyper-segmentation of the nucleus—a feature of neutrophil maturation (**Figure 5C**). High Ly6C expression, as seen in IFNAR^hi^ neutrophils, is associated with greater proinflammatory capacity in monocytes^49^. This suggests IFNAR^+^ neutrophils are a distinct, relatively mature BM subset that may also be more proinflammatory.

We tested whether IFN-I-sensing (i.e., IFNAR^+^) neutrophils extravasate into the meninges from the skull given: 1) our observation that CSD leads to increased neutrophil trafficking in vascular channels connecting skull BM to the meninges (**Figures 2G**-**I**), and 2) no evidence to suggest local meningeal proliferation (**Figure S6D**). Characterization of IFNAR^+^ neutrophils from several tissues revealed CSD-associated depletion was specific to skull BM (**Figure 5D**). Presumably this reflects egress from skull BM into the meninges, consistent with recent work from multiple labs showing skull BM provides a specialized reservoir for innate immune cell recruitment to the brain in conditions of neurological damage or disease^26–28^, and supported by our findings in cleared tissue. To our knowledge, this is the first evidence for such a phenomenon under conditions of psychosocial stress.

IFN-I depletion in C57BL/6J mice normalized CSD-related meningeal neutrophil levels (**Figure 6E**). Our attempts to replicate these effects in LysM*^gfp/+^* mice were less straightforward (see **Limitations**), though anhedonic behavior improved in the CSD+IFNAR treated mice (**Figure 6B**). Meningeal B cell levels were unaffected by anti-IFNAR treatment; specifically, CSD stress resulted in fewer B cells regardless of antibody treatment (**Figure 6F**). This would suggest B cell depletion precedes IFNAR-mediated neutrophil recruitment and is consistent with data from our time-course study in which meningeal B cell numbers decreased prior to an increase in meningeal neutrophils, i.e. by day 8 vs day 14 (**Figures 1H**, **S2C**). However, at present we also cannot exclude the possibility that B cell depletion and neutrophilia are completely independent events.

What processes drive B cell egress from, and neutrophil infiltration toward, the meninges following CSD stress? Whereas the mechanisms driving B cell depletion remain unclear, our previous work may shed light on the latter question. For example, we have shown that CSD stress leads to randomly occurring microhemorrhages in the neurovasculature^22,50^. As neutrophils are typically ‘first on the scene’ to sites of brain injury^51^, their active recruitment may facilitate progression, maintenance, or resolution of this stress-induced damage. Consistently, skull BM-to-meninges neutrophil trafficking has been demonstrated in hemorrhagic stroke, wherein blood brain barrier (BBB) damage is present^52^—though, notably, we find no evidence for infiltration of neutrophils or other leukocytes into the brain parenchyma of CSD-stressed mice^22,23^.

Another possibility is that chronic release of neutrophils into the bloodstream via repeated exposure to stress^15,16^ drives formation of these microhemorrhages. For example, in neurological conditions, neutrophils are notorious drivers of bystander damage due to their release of toxic reactive oxygen species (ROS)^51–53^. Our GSEA results (“GO: detoxification”, “GO: cell redox homeostasis”; **Figure S8**) prompted us to explore ROS production in blood and meningeal neutrophils. We saw no differences in ROS production between HC and CSD neutrophils on a per-cell basis (**Figure S11**), though the overall increase in ROS-producing neutrophils may exacerbate risk for bystander damage.

Blood-derived CSD neutrophils also showed signs of being both more rigid and proinflammatory (**Figure S9E**). We observed increased phalloidin staining intensity in CSD blood neutrophils (**Figure S9E**), consistent with enhanced formation of filamentous actin (F-actin)^54^. Increased F-actin formation may cause neutrophil rigidity, thereby increasing the likelihood of neutrophil ‘stalling’ in narrow brain parenchymal capillaries. Indeed, we saw more GFP^+^ neutrophils ‘stuck’ throughout the neurovasculature in CSD brains, and more neutrophil-like cells coprecipitating with neurovascular endothelia (NVE), despite aggressive exsanguination and tissue perfusion (**Figure S4**), consistent with another report^55^. Interestingly, in mouse models of Alzheimer’s disease (AD), neutrophil stalling was stochastic, resulting in reduced blood flow to the brain and associated memory deficits^56^. Our observations of NVE damage were similarly random and could possibly stem from stalling-related bystander damage, e.g., through neutrophil-mediated ROS-release.

Importantly, depressive symptoms are common in neurological disorders like stroke^57^ and AD^58^. This raises the possibility that altered neutrophil properties are a shared biological feature between neurological and psychiatric disorders; comparison of neutrophil phenotypes across brain disorders may provide fruitful insights into pharmacologically relevant treatment targets. In particular, we have identified IFN-I signaling as a putative mediator of meningeal neutrophil recruitment, which may worsen depressive-like symptoms (**Figure 6**). IFN-I signaling deleteriously impacts both neuronal injury after stroke^59^ and synapse loss in AD^60^. Given the relationship between depressive-like behavior and IFN-I^5,6,36^, and epidemiological evidence that depression doubles the risk for dementia late in life^61^, future work in this area is warranted.

## Limitations and strengths

While the social defeat paradigm is well-established and pharmacologically validated for the preclinical study of depression- and anxiety-like behavior^62^, a major limitation of this and similar studies is that they are done almost exclusively in adult male mice, though depression and anxiety disproportionately affect cis women^63^ and transgender youth^64^. ‘Resident’ CD-1 males commonly used in defeat paradigms to induce depressive-like behavior do not show aggression towards ‘intruder’ females unless manipulated to do so, either through surgical implant^65^ or by daily application of male odorant to females^66^, and unfortunately these adaptations to the standard paradigm were not published until after we had begun data collection.

Another limitation of our study is that social defeat stress is obligatorily associated with some degree of fight-inflicted wounding. We mitigated this by regularly clipping the CD-1 aggressor’s teeth, terminating defeat encounters before the 5 min duration of the stressor if an aggressive encounter involved a visible degree of biting, and removing CD-1 aggressors as stimulus animals if they were consistently hyper-aggressive. We also shaved mice at the time of sacrifice to assess severity of wounds. We note that without this step, the degree of wounding is not necessarily obvious under the fur. Anecdotally, higher wound scores were associated with greater levels of circulating and meningeal monocytes. This may require a more nuanced approach when attempting to translate results from social defeat stress models into a clinical setting, as has been discussed elsewhere^67^.

Finally, depletion of IFN-I signaling showed pleiotropic effects that were strain-dependent. We cannot account for the variation seen in LysM*^gfp/+^* CSD+IFNAR mice, though close inspection of HC data from different experiments suggests some individuals may be sensitive to the effects of repeated injection stress (**Figure S15D**). Our data also cannot rule out a role for IFNAR signaling in other cell types in behavior; future experiments using neutrophil-specific IFNAR knockouts will be important.

Our strengths include our whole organism approach to neutrophil dynamics. Triangulation and replication of our findings via multiple experimental approaches yielded strong evidence for chronic stress-induced meningeal neutrophilia. Moreover, data-led investigation of candidate mechanisms revealed IFN-I-mediated migration of neutrophils from skull BM to the meninges as a future target for translational study of stress-associated depression.

## Methods

### Animals

Strains used were either C57BL/6J male mice purchased from Jackson Labs (Bar Harbor, ME) or male LysM^+/*gfp*^ offspring from C57BL/6J mice bred in our facility with LysM*^gfp^*^/*gfp*^ mice, which strongly express GFP in neutrophils^68^ (obtained from Dr. Dorian McGavern, NINDS). Upon arrival into the facility, purchased mice were randomly pair-housed in divided cages and given one week of acclimation to the facility. Mice bred in-house were weaned at 3 weeks of age into same-sex cages of littermates. Sex was assigned based on external anatomy. Animals were housed under a reverse light cycle (lights off 8:00 AM to 8:00 PM), with food and water *ad libitum*. At 8-10 weeks of age, mice were randomly assigned to an experimental group. All procedures were approved by the National Institute of Mental Health Institutional Care and Use Committee and conducted in accordance with National Institutes of Health guidelines.

### Chronic social defeat (CSD)

CSD stress was performed as previously reported^22^. Briefly, the ‘intruder’ test mouse was introduced into the home cage of an aggressive, CD-1 (Taconic; Rensselaer, New York) retired breeder. The two were separated by a perforated barrier and given 24 h to acclimate; the barrier allowed for olfactory, visual, and auditory communication, but not tactile contact. Each day for either 1, 2, 4, 8, or 14 consecutive days, depending on the experiment, the barrier was lifted, and agonistic encounters were allowed to occur for 5 m. Interactions were monitored by a trained individual to ensure the test mouse exhibited submissive behavior and conversely that the CD1 exhibited dominant behavior. Efforts to minimize physical damage were taken, i.e. CD1 mice were lightly anesthetized with isoflurane and incisors were trimmed prior to starting the social defeat paradigm, and on a weekly basis thereafter. Test mice were shaved and inspected for the presence of wounds at the end of the experiment; wounds were scored on a scale from 1-10 (1 = no injuries, 5 = combination of old and new bite marks, 10 = severe wounds). Animals with wound scores of 10 were excluded.

### Behavioral phenotyping

All behavioral testing was done by both male and female experimenters during the dark phase of the light cycle, prior to a defeat encounter on that day. On the day of testing, mice were moved to a separate behavioral room with red lighting and acclimated for one hour prior to running the behavioral assays. Tests were usually run on separate days, but when multiple tests were run on a single day, the animals were given an hour to recover in between tests. Behavioral tests were performed in the following order:

### Urine scent marking (USM)

As described previously^24^. Briefly, mice were placed into a novel arena (50 x 50 x 50 cm) that contained a thick sheet of paper; one corner of this paper was ‘spotted’ with 50 μL of urine from multiple estrus females, and testing was performed with the lights off while the experimenter was outside the room. After testing, the sheets of paper were sprayed with ninhydrin and heated to indicate the presence of proteins, allowing for visualization of urine marking. Photos were taken of the sheets by an experimenter blind to group identity, then analyzed in ImageJ^69^. Reduced preference marking for the female scent is indicative of social anhedonia.

### Open field (OF)

Exploration of the novel open field arena, 50 x 50 x 50 cm in dimension, was performed under dim white lighting (∼25 lux). Mice were placed in the middle of the arena; total distance moved, crosses to center, and time in center over a 10 min testing period whilst the experimenter was out of the room were later analyzed with automated tracking software (Clever Sys TopScan Suite) to eliminate potential human bias. Fewer crosses to center, reduced time in center, and reduced movement are all indicative of increased anxiety-like behavior.

### Tissue collection

For all studies, mice were euthanized 2 h after their final exposure to the defeat stressor. Tissue collection occurred between ∼8am and noon, with modifications as indicated below. Venous blood was collected into EDTA tubes via puncture of the submandibular vein and kept on ice until processing. Retro-orbital injections were administered while mice were under light isoflurane anesthesia. After, mice were deeply anesthetized with isoflurane prior to perfusion with 35 mL room temperature PBS. When indicated, hindlegs were collected for tibial BM extraction. The head was decapitated, and intact skull cleaned with a scalpel to remove muscle and connective tissue.

### Histology

LysM*^+/gfp^* mice received a retro-orbital intravascular injection of either DyLite 649- or DyLite 594-conjugated tomato lectin (TomL, Cat #DL-1178, Vector Labs), which was allowed to circulate for 5 min to label blood vessels before perfusion. In addition to PBS, mice were perfused with 10 mL of 4% cold paraformaldehyde. Skulls were post-fixed in 4% PFA for 24 h at 4°C before transferring to 25% sucrose solution for cryoprotection, dehydration, and preparation for imaging.

### Single cell suspensions

WT mice were retro-orbitally injected with 4 μg CD45-FITC (Cat. #103108; Biolegend). Mice were then allowed to recover; after 25 min of circulation, mice were anesthetized lightly for venous blood collection. The decapitated skull was placed in cold HBSS + 0.1% BSA on ice until further processing.

### Tissue clearing and analysis of skull-to-meningeal vascular channels

We used the CUBIC tissue clearing method^70^ on whole skulls from LysM*^+/gfp^* mice. Tissue was prepared as for histology except that samples were transferred to PBS instead of sucrose after 24h fixation. Decalcification of bone prior to tissue clearing was achieved by incubation in decalcification solution (10% EDTA, 15% imidazole) for 5-7 days at 37°C with shaking^71^. The decalcification solution was refreshed once on day 3. Following tissue clearing, the whole skull was inverted and placed in a glass-bottom dish filled with fresh Reagent 2^70^ and imaged using a Zeiss 780 confocal microscope fitted with 10x objective. 2-5 images were collected at random locations for each skull and analyzed using IMARIS 9.7. For each image, both the number of vascular channels (labeled with TomL) and the number of discreet neutrophils (GFP^+^) in a channel were counted. For each individual sample, the ratio of averaged neutrophils normalized to the average number of channels is presented.

### Peripheral blood preparations

∼500 μL of venous blood was lysed in 8 mL ACK Lysis Buffer (Cat. # 351-029-721; Quality Biological, Inc.) for 5 min at room temperature, and the reaction stopped by diluting with 7 mL cold HBSS + 0.1% BSA. Cells were pelleted, washed, and prepared for staining.

### Skull and tibial BM preparations

To prepare for skull BM extraction, the dorsal calvarium was trimmed to be relatively flat, and meninges were removed under a dissecting microscope. Next, the skull was cut into small bone pieces with scissors in cold HBSS + 0.1% BSA. This entire slurry was transferred to a 70 μm cell strainer and mashed with the rubber end of a 3 mL syringe for approximately 2 min per sample. Tibia were prepared by first stripping away all tissue from the bone, then cutting the very top such that a 23g syringe needle could be inserted to flush out the BM into a tube of cold HBSS + 0.1% BSA. This was next transferred to a 70 μm cell strainer and mashed with the rubber end of a 3 mL syringe. For both kinds of BM, the resulting cell suspensions were then pelleted and prepared for flow cytometry.

### Meningeal dissection and single cell suspension

Meninges samples were collected by first cutting around the lateral sutures of the skull; the dorsal skull and ventral skull were transferred to a fresh petri dish filled with cold HBSS + 0.1% BSA and kept on ice while pial and arachnoid meningeal membranes were gently picked off the entire outer surface of the brain into a second ‘working dish’ with Dumont #5 forceps (Cat. #RS-5058; Roboz, Gaithersburg, MD). Extra care was taken to avoid inclusion of choroid plexus from the 4^th^ ventricle. Once finished with the brain, skull pieces were transferred as necessary into the working dish to remove attached meninges; we avoided leaving skull pieces in the ‘working dish’ to minimize contamination with cells from skull BM. Upon completion of the meningeal dissection, samples were transferred to a fresh tube and cells were pelleted by centrifugation, then resuspended in 2mL of BSA-free HBSS supplemented with 2.5 mg/mL Collagenase D (Cat. #11088858001; Roche) and 12.5 μL of 0.5 mg/mL DNAseI (Cat. #L5002139; Worthington) for cell dissociation. The samples were incubated at 37°C for 30 min, diluted with cold HBSS + 0.1% BSA, and mashed through a 70 μm cell strainer into single cell suspension.

### Meningeal whole mount preparations and staining

The dorsal skull was carefully removed to retain maximal attachment of the meningeal layers from tissue prepared for histology, and brains were returned to fresh 25% sucrose until sunk for further staining (see **Supplemental Methods**). Dorsal meninges were gently peeled from the skull as a single sheet, mounted ventral (brain) side up/dorsal (skull) side down onto slides and encircled with a Pap pen.

### Meningeal immunohistochemistry

Meningeal whole mount samples from LysM*^+/gfp^* mice were dried, washed with PBS, blocked for 1 h in 4% normal goat serum in 0.4% Triton-PBS, and incubated in a humidity chamber overnight at room temperature with chicken anti-GFP (1:1000, Cat #13970, Abcam), diluted in 0.2% Triton-PBS with 2% normal goat serum. Approximately 18h later, the samples were washed 3 x 5m with PBS, and incubated for 2 h with Chicken IgY-Alexa Fluor 488 (1:500, Cat #150169, Abcam) in 0.4% Triton-PBS. Samples were washed 2 x 5m with PBS, given a quick rinse in deionized water, and counterstained with DAPI for 5 minutes. They were rinsed briefly again with deionized water then cover-slipped with PVA-DABCO (made in-house).

### Image acquisition and analysis

Meningeal whole mounts were tile scanned using a 20x objective at 1024×1024 resolution and online stitching with a Zeiss 780 confocal microscope. Two independent investigators blind to treatment hand-counted confocal images of the meningeal whole mounts to quantify the density and location of neutrophils (characterized by high GFP expression, irregularly shaped nuclei, and a semi-round shape) within the tissue using ImageJ software^69^. There was excellent agreement in their counts (Pearson correlation, \*\*\*\**p* < 0.0001, *r* = 0.937). Neutrophils were also examined in relationship to blood vessels; neutrophils > 10 μm from a blood vessel were provisionally called ‘parenchymal,’ whereas neutrophils ≤ 10 μm were called ‘abluminal.’ Neutrophils were otherwise considered intravascular.

### Cell staining and flow cytometry

To exclude dead cells, samples were stained with either Fixable Viability Dye eFluor™ 780 (Cat. #65-0865; eBioscience) for 10 min at room temperature at 1:2400 dilution, or with Zombie AQUA (Cat # 423102, Biolegend) for 15 min at room temperature at 1:100 dilution. The cells were then washed and blocked with 1 μL of normal goat serum (Cat #G9023-10ML; Sigma) and 1 μL of Fc block (Cat #101302; Biolegend) for 10m on ice. 25 μL Brilliant Violet stain buffer (Cat #563794; BD Horizon) was then added, followed by an antibody master mix (**Table S7**). Cells were incubated on ice for 25 min, washed with PBS, and fixed in 2% PFA at room temperature for 15 min for analysis on either a BD LSR Fortessa or a Beckman Coulter CytoFLEX flow analyzer. Compensation was performed for each session using UltraComp eBeads (eBioscience 01-2222-42) conjugated to antibodies used in the sample panels. Viability dye and GFP^+^ controls used cells instead of beads. Data were analyzed using FlowJo (BD) software with manual gating. Absolute cell counts were determined using CountBrite counting beads (ThermoFisher, catalog #C36950). See **Supplemental Methods** for details on imaging flow cytometry.

### Meningeal scRNAseq

Meningeal scRNAseq data were acquired from 8 non-stressed HC mice and 4 stress-susceptible CSD mice as described previously^21^. In brief, live, nucleated, singlet cells (DAPI^-^DRAQ^+^) were sorted on a BD FACS Aria Fusion into HBSS + 10% FBS prior to droplet encapsulation using 10x Genomics’ Drop-seq platform (Chromium v2). 3 pooled samples of 4 mice each were generated (two HC pools, one CSD pool) and run on the same 10x Chromium chip; libraries were sequenced on Illumina NextSeq 550 and feature counts generated using Cellranger V2 pipeline. See **Supplemental Methods** for additional information.

### Cytospin

Skulls were obtained and prepared for single-cell suspension. The cells were then fixed, pelleted onto thin coverslips using funnel centrifugation (Cat #10-354, Fisher HealthCare), and imaged using a Zeiss 780 confocal microscope fitted with 10x objective.

### IFNAR-blocking assay

Anti-IFNAR (clone: MAR1-5A3, Cat # BE0241) and non-specific, IgG isotype control (clone: MOPC-21, Cat # BE0083) antibodies for repeated *in vivo* injections were purchased from BioXCell and diluted in *InVivo*Pure pH 7.0 Dilution Buffer (Cat # IP0070) to a concentration of 5 mg/mL. Mice were i.p. injected with 1mg of antibody on day zero (d0), before the start of the defeat paradigm, and received 0.5 mg of antibody every third day thereafter, for a total of 5 injections per mouse. Defeats were done approximately 1-3 h after injections. LysM^+/*gfp*^ mice were ear-tagged and randomly assigned to one of three groups: HC+IgG, CSD+IgG, or CSD+IFNAR. WT mice were ear-tagged and randomly assigned to one of four groups: HC+IgG, HC+IFNAR, CSD+IgG, or CSD+IFNAR.

### Statistics

Prism 9.0.2 (GraphPad Software, LLC) was used for statistical testing and graphing of univariate analyses. Normality and equal variance were assessed, and appropriate tests were conducted thereafter. For simple, two-group comparisons, samples with non-equal variance were analyzed using a Mann-Whitney U test, whereas samples with equal variance were analyzed using a Student’s t test. In almost all instances CSD-stressed mice failed to mark in the USM test; the behavioral output was thus better modeled statistically as a binary response (did or did not mark). We therefore used χ^2^ tests for univariate analyses of group in the USM test; posthoc testing was performed with the chisq.posthoc.test package in R, using the Benjamini-Hochberg method for multiple comparisons corrections. For multiple comparisons of normally distributed data, ordinary one- or two-way ANOVAs were used for analysis. When a main effect was present, post-hoc analyses were conducted (Dunnett’s or Tukey’s, respectively). For multiple comparisons of non-parametric data, a Kruskal-Wallis test was used, and Dunn’s test for multiple corrections was run for post-hoc analysis. Univariate comparisons are summarized as the mean ± SEM and considered statistically significant at *p* < 0.05. No power calculations were performed at the outset of the experiment. Please see **Supplemental Methods** for more information about statistical modeling.

## Data availability

Preprocessed single cell RNA sequencing data can be accessed at https://zenodo.org/records/13378961. Microarray data have been deposited in GEO under accession code GSE275966. Additional supporting data is available upon request.

## Code availability

All code used for analysis is available at https://github.com/staceykigar/meningeal_neut.

## Competing interests

E.T.B. is a consultant for Sosei Heptares. M.L.L. is currently employed at AstraZeneca but was an employee at NIH at the time this work was conducted. The other authors have no conflicts to declare.

## Acknowledgements

We thank Dorian McGavern for helpful commentary on early versions of the manuscript and for kindly providing LysM*^gfp/gfp^* mice. We thank Monica Manglani, Hannah Mason, Methma Udawatta, Donghyun Kim, and Chris Higham for technical assistance. We thank staff in the Laboratory of Genome Integrity (Flow Cytometry Core, National Cancer Institute), staff in the Microarrays and Single-Cell Genomics Core (National Human Genome Research Institute), and veterinary care staff for their help with data collection.

This work was supported by the NIMH Intramural Research Program, ZIA MH001090 (S.L.K., A.E.D., R.A., V.H.S., J.D.S., N.E.E., C.N.P., M.L.L., S.J.L, M.H.); the NIHR Cambridge Biomedical Research Centre (S.L.K.); and the UK Medical Research Council, award MR/S006257/1 (M.E.L.). All research at the Department of Psychiatry in the University of Cambridge is supported by the NIHR Cambridge Biomedical Research Centre (NIHR203312) and the NIHR Applied Research Collaboration East of England. The views expressed are those of the author(s) and not necessarily those of the NIHR or the Department of Health and Social Care.

## Author Contributions

S.L.K., A.E.D., R.A., J.D.S., N.E.E., C.N.P., M.L.L., S.J.L., F.L., A.G.E. participated in data collection. S.L.K., M.E.L., V.H.S., F.L., A.G.E. performed analyses. S.L.K. and M.E.L. conceived the study and wrote the manuscript. M.R.C. and E.T.B. contributed expertise. A.E.D., R.A., J.D.S., V.H.S., C.N.P., M.L.L., S.J.L., F.L., M.R.C., E.T.B., M.H. reviewed and edited the manuscript.

## Notes

https://zenodo.org/records/13378961

https://www.ncbi.nlm.nih.gov/geo/query/acc.cgi?acc=GSE275966

https://github.com/staceykigar/meningeal_neut

