## Supplemental Materials for "Chronic social defeat stress induces meningeal neutrophilia via type I interferon signaling"

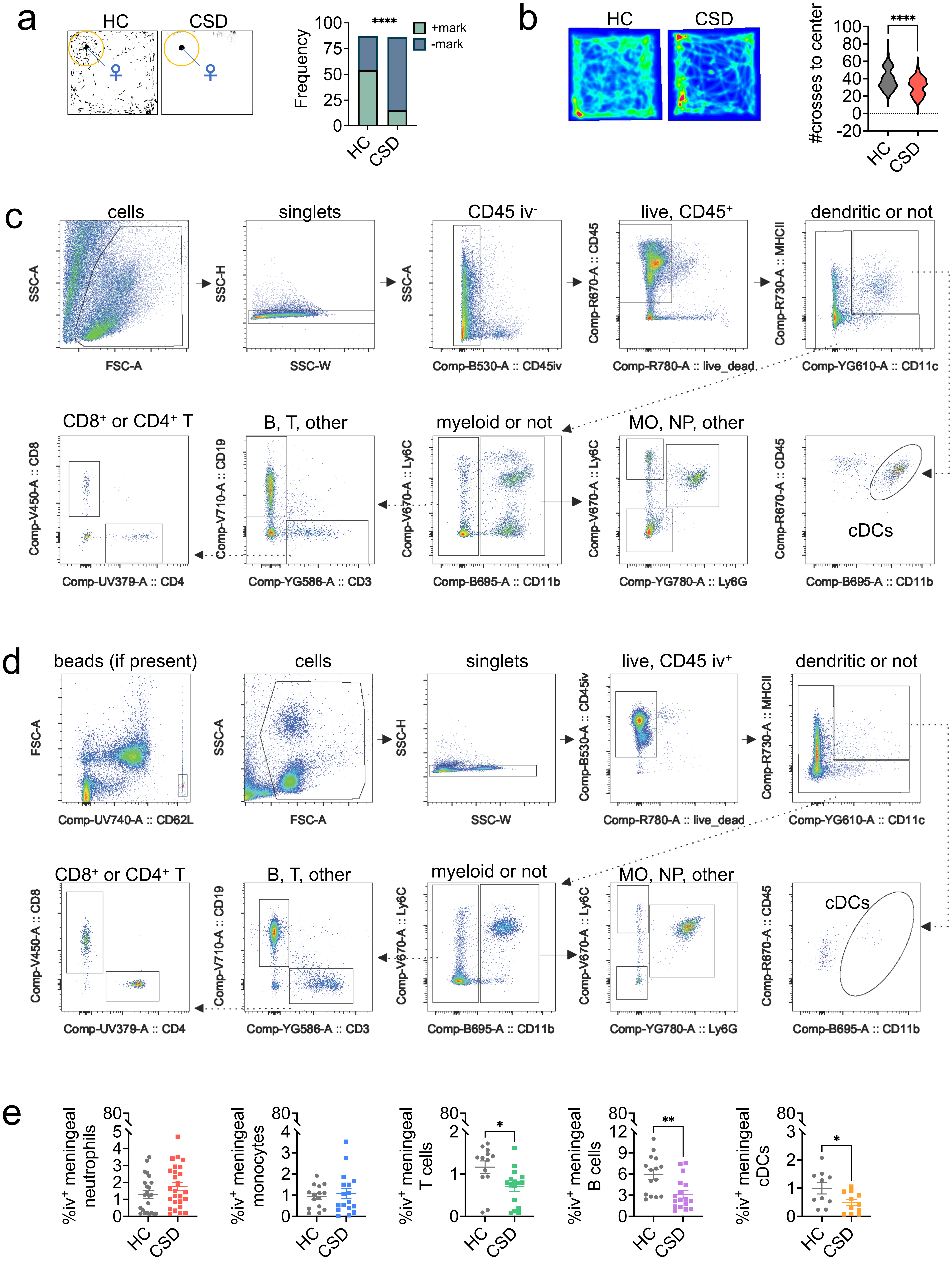

a
HC
CSD
Frequency
100
75
50
25
0
****
+mark
-mark
b
HC
CSD
#crosses to center
****
c
cells
singlets
CD45 iv⁻
live, CD45⁺
dendritic or not
SSC-A
FSC-A
SSC-H
SSC-W
Comp-B530-A :: CD45iv
Comp-R670-A :: CD45
Comp-R780-A :: live_dead
Comp-R730-A :: MHCII
Comp-YG610-A :: CD11c
CD8⁺ or CD4⁺ T
B, T, other
myeloid or not
MO, NP, other
cDCs
Comp-V450-A :: CD8
Comp-UV379-A :: CD4
Comp-V710-A :: CD19
Comp-YG586-A :: CD3
Comp-V670-A :: Ly6C
Comp-B695-A :: CD11b
Comp-YG780-A :: Ly6G
d
beads (if present)
cells
singlets
live, CD45 iv⁺
dendritic or not
FSC-A
Comp-UV740-A :: CD62L
Comp-B530-A :: CD45iv
e
%iv⁺ meningeal neutrophils
%iv⁺ meningeal monocytes
%iv⁺ meningeal T cells
%iv⁺ meningeal B cells
%iv⁺ meningeal cDCs
*
**


Figure S1

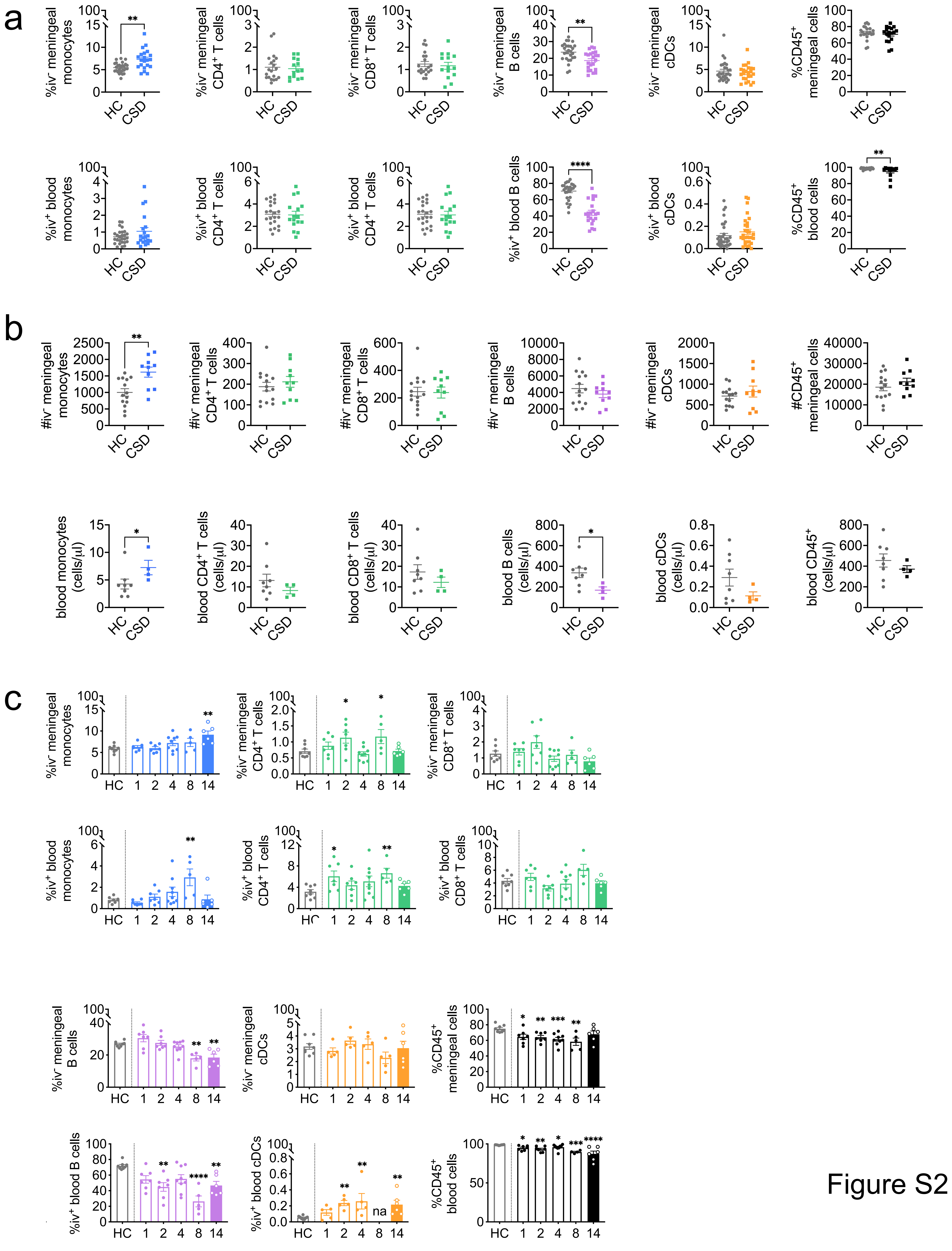


Figure S2

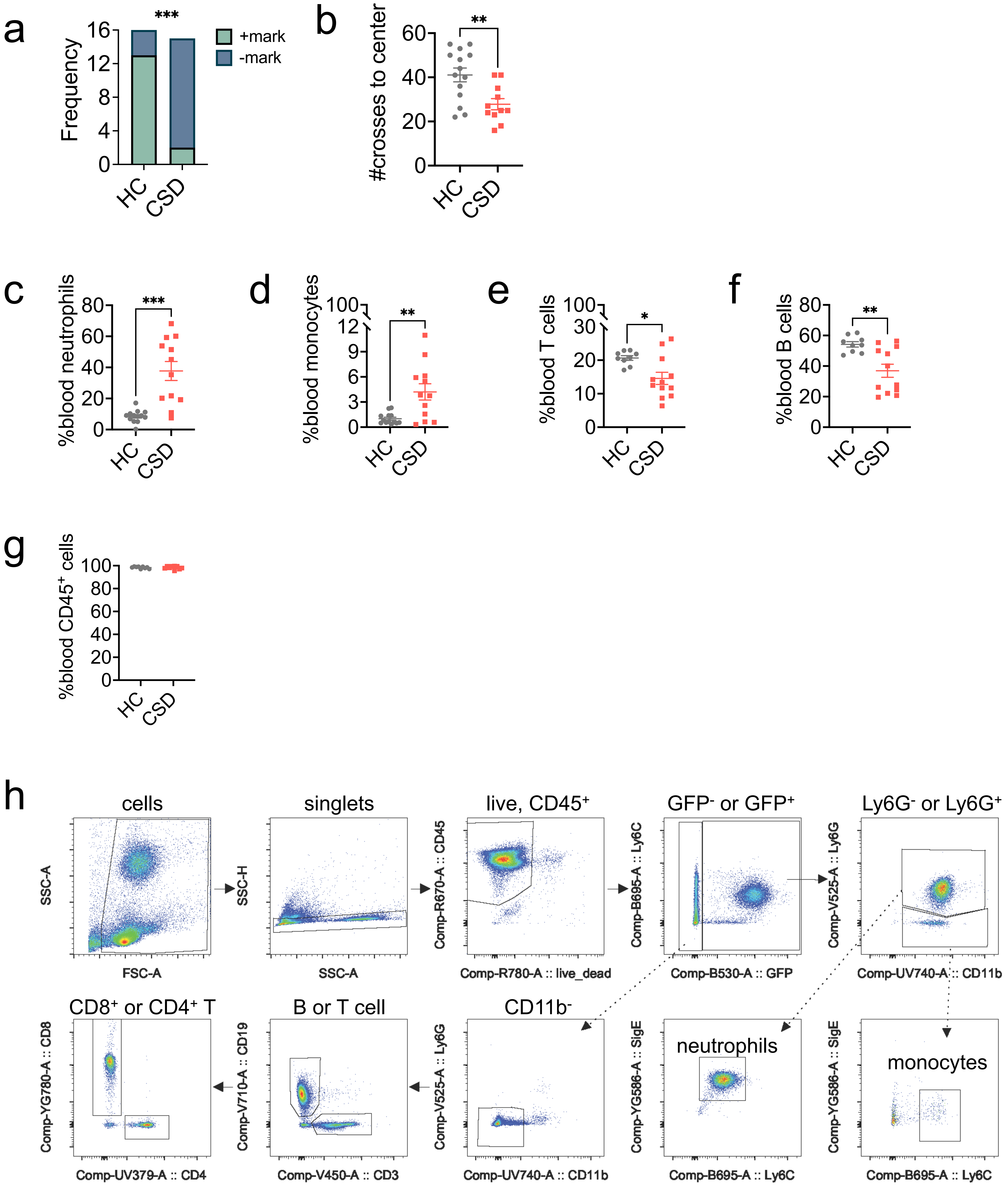

a
***
+mark
-mark
Frequency
16
12
8
4
0
HC
CSD
b
**
#crosses to center
60
40
20
0
HC
CSD
c
***
%blood neutrophils
80
60
40
20
0
HC
CSD
d
**
%blood monocytes
100
12
9
6
3
0
HC
CSD
e
*
%blood T cells
100
30
20
10
0
HC
CSD
f
**
%blood B cells
80
60
40
20
0
HC
CSD
g
%blood CD45+ cells
100
80
60
40
20
0
HC
CSD
h
cells
SSC-A
FSC-A
singlets
SSC-H
SSC-A
live, CD45+
Comp-R670-A :: CD45
Comp-R780-A :: live_dead
GFP- or GFP+
Comp-B695-A :: Ly6C
Comp-B530-A :: GFP
Ly6G- or Ly6G+
Comp-V525-A :: Ly6G
Comp-UV740-A :: CD11b
CD8+ or CD4+ T
Comp-YG780-A :: CD8
Comp-UV379-A :: CD4
B or T cell
Comp-V710-A :: CD19
Comp-V450-A :: CD3
CD11b-
Comp-V525-A :: Ly6G
Comp-UV740-A :: CD11b
neutrophils
Comp-YG586-A :: SigE
Comp-B695-A :: Ly6C
monocytes
Comp-YG586-A :: SigE
Comp-B695-A :: Ly6C


Figure S3

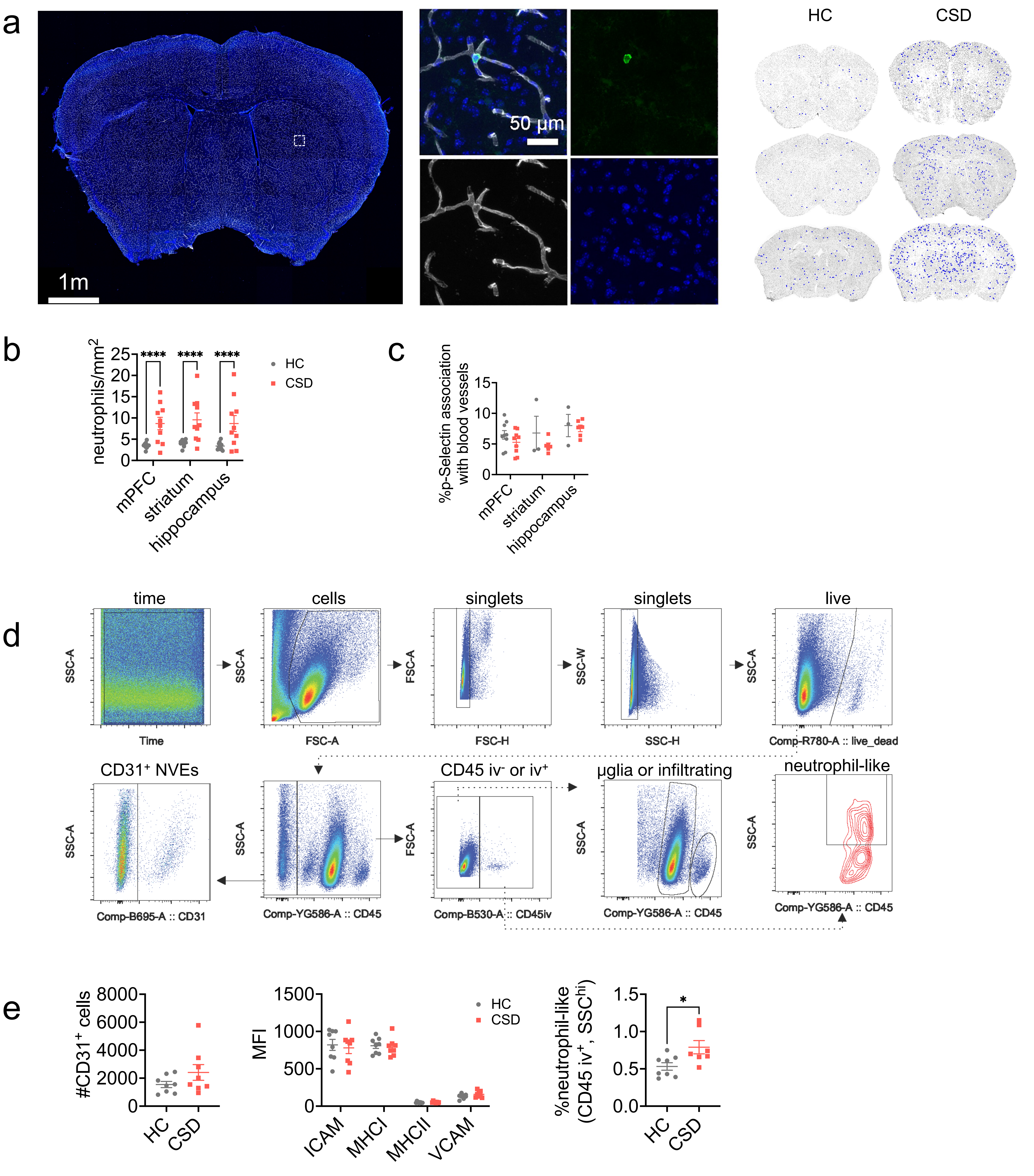
a
1m
50 µm
HC
CSD
b
neutrophils/mm²
****
HC
CSD
mPFC
striatum
hippocampus
c
%p-Selectin association with blood vessels
mPFC
striatum
hippocampus
d
time
cells
singlets
singlets
live
SSC-A
Time
FSC-A
FSC-H
SSC-W
SSC-H
Comp-R780-A :: live_dead
CD31⁺ NVEs
CD45 iv⁻ or iv⁺
µglia or infiltrating
neutrophil-like
Comp-B695-A :: CD31
Comp-YG586-A :: CD45
Comp-B530-A :: CD45iv
e
#CD31⁺ cells
HC
CSD
MFI
ICAM
MHCI
MHCII
VCAM
%neutrophil-like (CD45 iv⁺, SSC^hi)
*

Figure S4

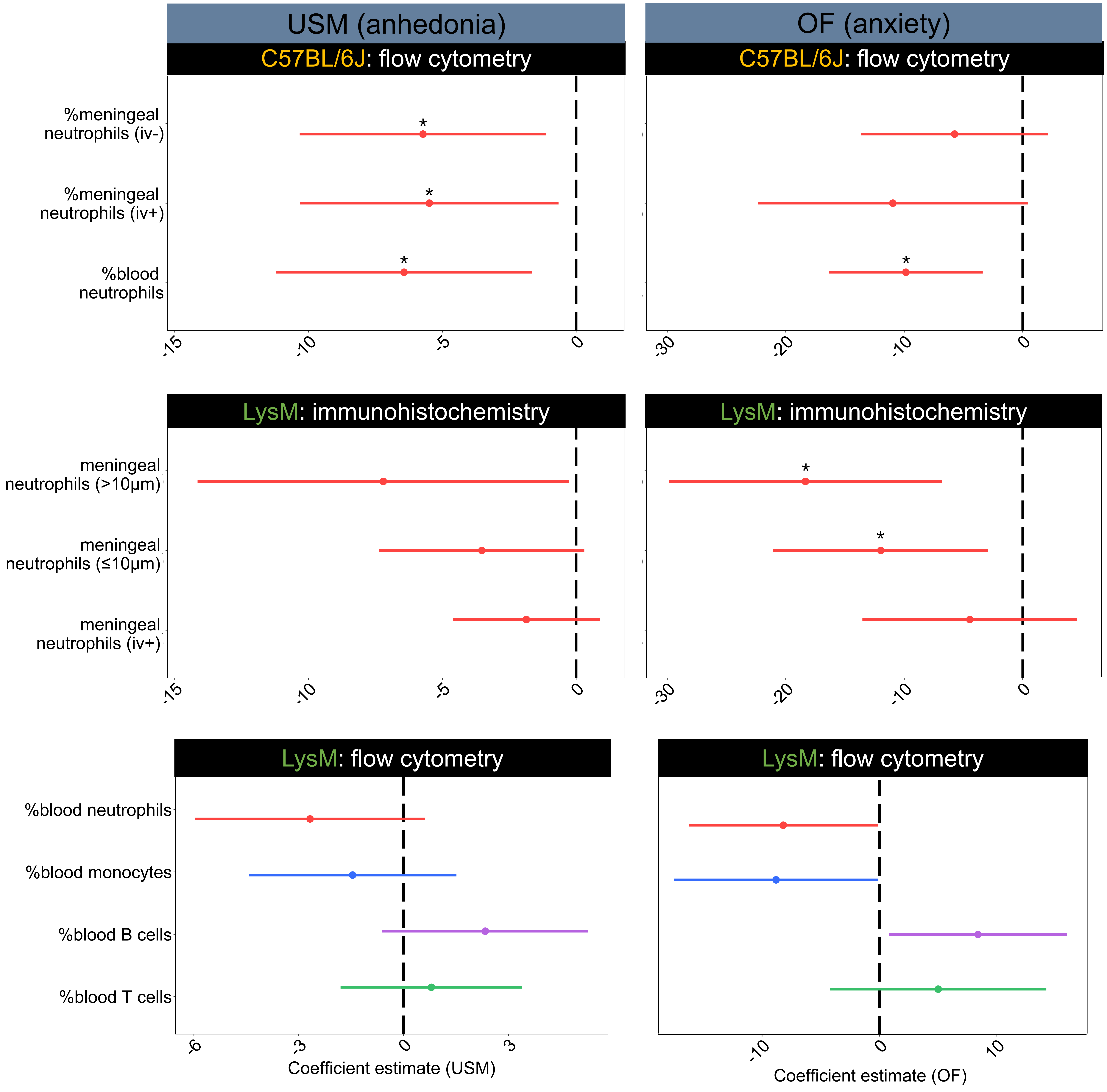

USM (anhedonia)
OF (anxiety)
C57BL/6J: flow cytometry
C57BL/6J: flow cytometry
%meningeal neutrophils (iv-)
%meningeal neutrophils (iv+)
%blood neutrophils
*
*
*
*
-15
-10
-5
0
-30
-20
-10
0
LysM: immunohistochemistry
LysM: immunohistochemistry
meningeal neutrophils (>10μm)
meningeal neutrophils (≤10μm)
meningeal neutrophils (iv+)
*
*
-15
-10
-5
0
-30
-20
-10
0
LysM: flow cytometry
LysM: flow cytometry
%blood neutrophils
%blood monocytes
%blood B cells
%blood T cells
-6
-3
0
3
Coefficient estimate (USM)
-10
0
10
Coefficient estimate (OF)


Figure S5

**a** Marker genes by % of cells expressing gene

| Microglia | Preneutrophils | Neutrophils | Monocytes Ly6Chi | Monocytes NOS | Border-associated macrophage | Perivascular macrophage (Cd206) | cDCs | pDCs | Mast cells and others | Endothelial and others | Fibroblasts and other | Olfactory neurons | Pro B cells and other precursors | Pre B cells | Mature B cells 1 | Mature B cells 2 | T cells 1 | T cells 2 | NK cells | Erythrocytes |
|---|---|---|---|---|---|---|---|---|---|---|---|---|---|---|---|---|---|---|---|---|
| P2ry12 | Gm10282 | Cxcr2 | Asf1b | Fn1 | Ms4a7 | Mrc1 | Cd209a | Ccr9 | Cdk6 | Crip2 | Col1a2 | Elavl3 | Myl4 | Arl5c | Ms4a1 | Fcmr | Cd3g | Cd3g | Klrb1c | Trim10 |
| Gpr34 | 1700020L24Rik | Hdc | Cdca3 | Ms4a8a | Cxcl16 | Dab2 | Tnip3 | Cox6a2 | Cpa3 | Krt18 | Rspo3 | Fstl5 | Mzb1 | Fcrla | Tnfrsf13c | Gm31243 | Il7r | Cd3d | Klre1 | Gypa |
| Siglech | Orm1 | Trem3 | Ccna2 | Clec4a1 | Slamf9 | Stab1 | Clec4b1 | Sh3bgr | Cst7 | Flt1 | Serpinh1 | Gnal | Gm30211 | Ebf1 | Siglecg | H2-DMb2 | Thy1 | Thy1 | Klrk1 | Slc4a1 |
| Tmem119 | Ms4a3 | Slc7a11 | Racgap1 | Clec4a3 | C3ar1 | Ms4a7 | H2-DMb1 | Cd300c | Gata2 | Igfbp7 | Aebp1 | Scn9a | Ezh2 | Tifa | Fcmr | Ms4a1 | Lat | Ms4a4b | Klrd1 | Alas2 |
| Olfml3 | Spc25 | Mmp9 | Mki67 | Ccr2 | H2-DMb1 | Cbr2 | Cbfa2t3 | Siglech | Ms4a2 | Timp3 | Cald1 | Sult1d1 | Bcl7a | Vpreb3 | Cd72 | Bank1 | Cd3d | Lck | Xcl1 | Tspo2 |
| Fcrls | Ccnb2 | Fpr2 | Smc2 | Ms4a4c | Ctsc | Pf4 | Slamf7 | Klk1 | Csrp3 | Hspb1 | Pcolce | Clgn | Smarca4 | Pafah1b3 | Spib | Cd79a | Skap1 | Gimap4 | Ncr1 | Cldn13 |
| Crybb1 | Cdkn3 | Pilra | Knstrn | F13a1 | Tmem176a | Maf | Ccr2 | Cd7 | Srm | Ly6c1 | Efemp1 | Snap25 | Sox4 | Cd72 | Cd79a | Ebf1 | Cxcr6 | Skap1 | Txk | Rhd |
| Mafb | Inhba | Mxd1 | Tacc3 | AI839979 | Lgmn | Folr2 | Ccnd1 | Upb1 | Sfxn1 | Tcf4 | Plpp3 | Flrt1 | Lockd | Siglecg | Ebf1 | H2-Ob | Bcl11b | Cd8b1 | Ctsw | Fech |
| Hpgds | Cdca8 | Il1r2 | Ccnb2 | Mcub | Tmem176b | Gas6 | Mgl2 | Runx2 | Rcl1 | Krt8 | Igfbp5 | Ttll7 | Igll1 | Sox4 | Vpreb3 | Cd79b | H2-Q7 | Gimap3 | Cd7 | Sox6 |
| Ecscr | Mki67 | Chil1 | Spc24 | Cd300a | Aif1 | Lyve1 | Pid1 | P2ry14 | Gnl3 | Fxyd6 | Itm2a | Pcolce2 | Akap12 | Cnp | Pax5 | Gm8369 | Ccnd2 | Ctsw | Ms4a4b | Snca |

**b** Marker genes by level of expression

| Microglia | Preneutrophils | Neutrophils | Monocytes Ly6Chi | Monocytes NOS | Border-associated macrophage | Perivascular macrophage (Cd206) | cDCs | pDCs | Mast cells and others | Endothelial and others | Fibroblasts and other | Olfactory neurons | Pro B cells and other precursors | Pre B cells | Mature B cells 1 | Mature B cells 2 | T cells 1 | T cells 2 | NK cells | Erythrocytes |
|---|---|---|---|---|---|---|---|---|---|---|---|---|---|---|---|---|---|---|---|---|
| Cst3 | Camp | S100a8 | Pclaf | Lyz2 | Cd74 | Apoe | H2-DMa | Irf8 | Rpl15 | Crip2 | Malat1 | Calm1 | Ptma | Vpreb3 | Ly6d | Rpl18a | Rpl17 | Cd3d | Nkg7 | Hba-a1 |
| Hexb | Trem3 | Retnlg | Lgals1 | S100a4 | H2-Aa | C1qa | H2-DMb1 | Rpl31 | Rps12 | Ly6a | Mgp | Omp | Ptprcap | Ebf1 | Cd79a | Rps19 | Rps14 | Rps15a | AW112010 | Hbb-bs |
| Lgmn | H2afz | Cxcr2 | S100a10 | Lgals3 | H2-Eb1 | Pf4 | Rps11 | Plac8 | Cmtm7 | Krt18 | Plpp3 | Gng13 | Stmn1 | Chchd10 | Cd79b | Fcmr | Tpt1 | Hcst | Ccl5 | Hbb-bt |
| P2ry12 | Ngp | Bmx | Ly6c2 | Ifitm3 | H2-Ab1 | Selenop | Plbd1 | Bst2 | Rpl14 | Hspb1 | Serpinf1 | Nsg1 | Tubb5 | Arl5c | Ms4a1 | Rps27 | Rplp1 | Cd3g | Klrk1 | Car2 |
| Tmem119 | Hmgb2 | S100a9 | Tmsb10 | F13a1 | Tmem176b | Ctsb | Gm2a | Rpl10 | Srgn | Ly6c1 | Id3 | Stoml3 | Hmgb1 | Dnajc7 | Siglecg | Ltb | Il7r | Rps16 | Klrb1c | Gpx1 |
| C1qc | Hmgn2 | Cxcl2 | Tuba1b | Ms4a6c | Aif1 | Dab2 | Rps9 | Siglech | Ifitm1 | Tm4sf1 | Igfbp5 | Map1b | H2afv | Tifa | Btg1 | Rpl13 | Rpl36a | B2m | Ncr1 | Hba-a2 |
| Ctss | Wfdc21 | Gsr | Pycard | Napsa | Ctsh | Cd68 | Tnip3 | Cox6a2 | Ctsg | Cmtm8 | 1500015O10Rik | Tuba1a | Mzb1 | Xrcc6 | Ifi30 | H2-DMb2 | Rpl19 | Gimap4 | Klrd1 | Prdx2 |
| Sparc | Pglyrp1 | Slc7a11 | Ppia | Fn1 | Tmem176a | Maf | Naaa | Sec61b | Npm1 | Flt1 | Col1a2 | Calm2 | Smarca4 | Pafah1b3 | Cd37 | Rps29 | Rplp0 | Rpl13a | Xcl1 | Gypa |
| Selplg | Lcn2 | Itgam | Ran | Psap | Slamf9 | Cbr2 | Cdkn1a | Rpl36al | Ier3 | Arhgap31 | Rbp1 | Tshz2 | Pgls | Rhoh | Tnfrsf13c | Gm31243 | Rps5 | Ms4a4b | Gzma | Cd24a |
| C1qb | Anxa1 | B430306N03Rik | Crip1 | Smpdl3a | Fth1 | Ftl1 | Cd209a | Tcf4 | Cst7 | Sparcl1 | Klf9 | Plekhb1 | Pkig | Fam53b | Cd72 | Bank1 | Cd163l1 | Rps13 | Id2 | Alas2 |

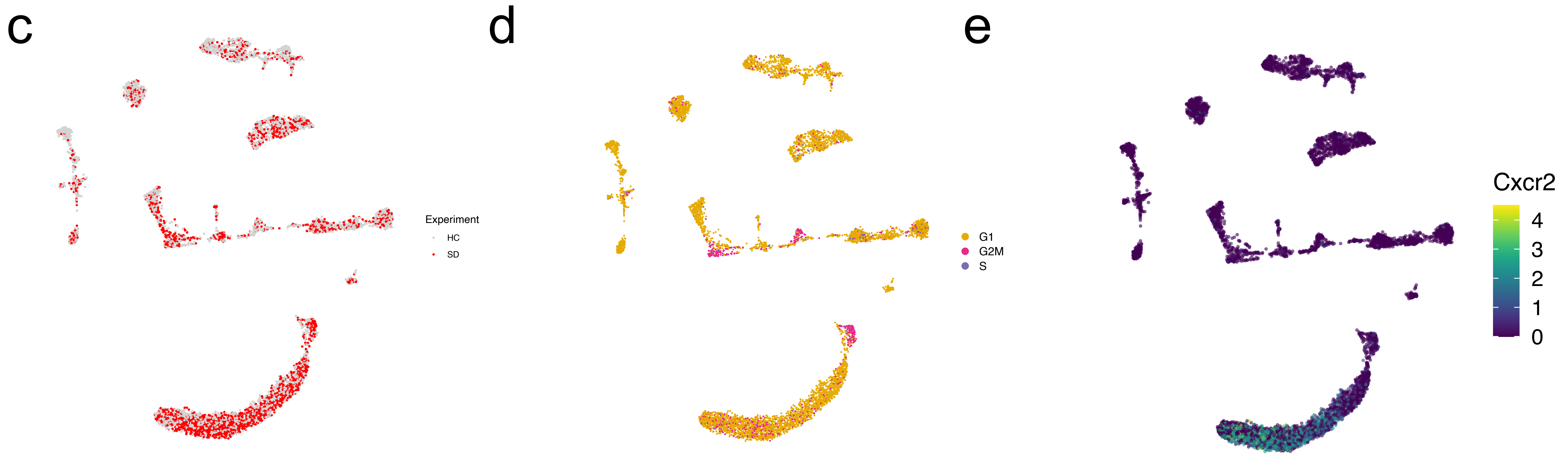


Figure S6

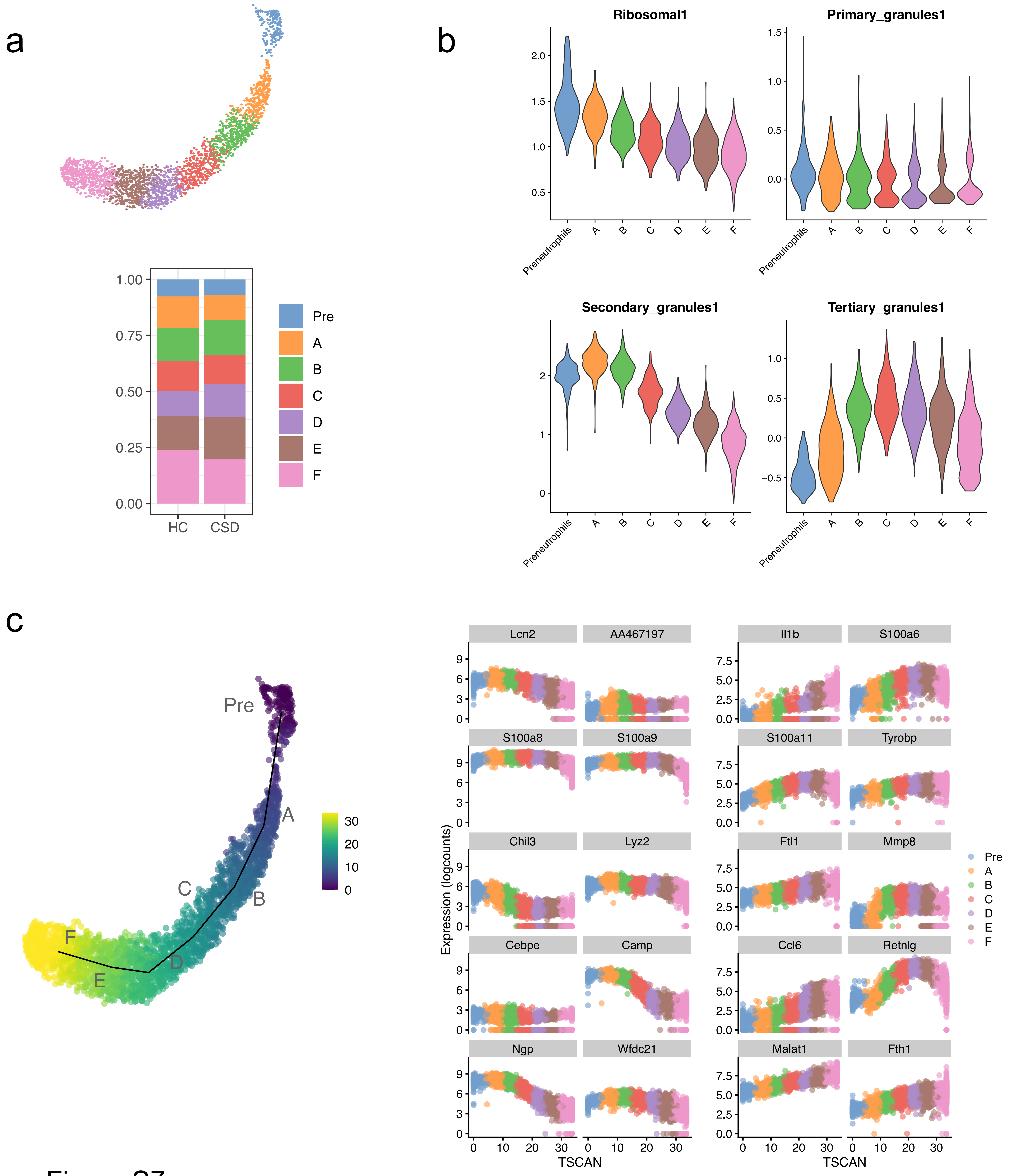

a
b
c
Pre
A
B
C
D
E
F
HC
CSD
Ribosomal1
Primary_granules1
Secondary_granules1
Tertiary_granules1
Preneutrophils
Lcn2
AA467197
Il1b
S100a6
S100a8
S100a9
S100a11
Tyrobp
Chil3
Lyz2
Ftl1
Mmp8
Cebpe
Camp
Ccl6
Retnlg
Ngp
Wfdc21
Malat1
Fth1
Expression (logcounts)
TSCAN


Figure S7

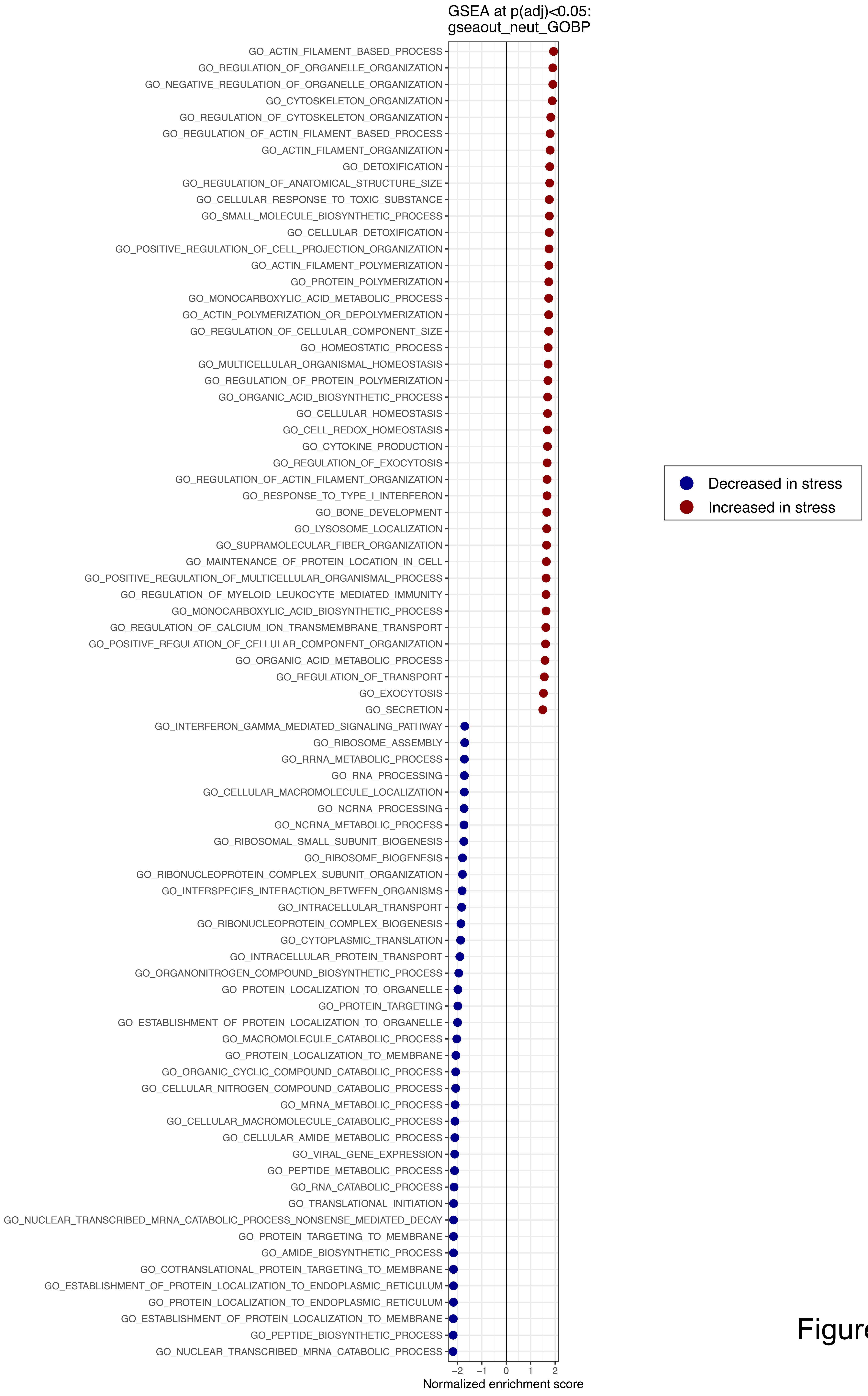
GSEA at p(adj)<0.05:
gseaout_neut_GOBP
GO_ACTIN_FILAMENT_BASED_PROCESS
GO_REGULATION_OF_ORGANELLE_ORGANIZATION
GO_NEGATIVE_REGULATION_OF_ORGANELLE_ORGANIZATION
GO_CYTOSKELETON_ORGANIZATION
GO_REGULATION_OF_CYTOSKELETON_ORGANIZATION
GO_REGULATION_OF_ACTIN_FILAMENT_BASED_PROCESS
GO_ACTIN_FILAMENT_ORGANIZATION
GO_DETOXIFICATION
GO_REGULATION_OF_ANATOMICAL_STRUCTURE_SIZE
GO_CELLULAR_RESPONSE_TO_TOXIC_SUBSTANCE
GO_SMALL_MOLECULE_BIOSYNTHETIC_PROCESS
GO_CELLULAR_DETOXIFICATION
GO_POSITIVE_REGULATION_OF_CELL_PROJECTION_ORGANIZATION
GO_ACTIN_FILAMENT_POLYMERIZATION
GO_PROTEIN_POLYMERIZATION
GO_MONOCARBOXYLIC_ACID_METABOLIC_PROCESS
GO_ACTIN_POLYMERIZATION_OR_DEPOLYMERIZATION
GO_REGULATION_OF_CELLULAR_COMPONENT_SIZE
GO_HOMEOSTATIC_PROCESS
GO_MULTICELLULAR_ORGANISMAL_HOMEOSTASIS
GO_REGULATION_OF_PROTEIN_POLYMERIZATION
GO_ORGANIC_ACID_BIOSYNTHETIC_PROCESS
GO_CELLULAR_HOMEOSTASIS
GO_CELL_REDOX_HOMEOSTASIS
GO_CYTOKINE_PRODUCTION
GO_REGULATION_OF_EXOCYTOSIS
GO_REGULATION_OF_ACTIN_FILAMENT_ORGANIZATION
GO_RESPONSE_TO_TYPE_I_INTERFERON
GO_BONE_DEVELOPMENT
GO_LYSOSOME_LOCALIZATION
GO_SUPRAMOLECULAR_FIBER_ORGANIZATION
GO_MAINTENANCE_OF_PROTEIN_LOCATION_IN_CELL
GO_POSITIVE_REGULATION_OF_MULTICELLULAR_ORGANISMAL_PROCESS
GO_REGULATION_OF_MYELOID_LEUKOCYTE_MEDIATED_IMMUNITY
GO_MONOCARBOXYLIC_ACID_BIOSYNTHETIC_PROCESS
GO_REGULATION_OF_CALCIUM_ION_TRANSMEMBRANE_TRANSPORT
GO_POSITIVE_REGULATION_OF_CELLULAR_COMPONENT_ORGANIZATION
GO_ORGANIC_ACID_METABOLIC_PROCESS
GO_REGULATION_OF_TRANSPORT
GO_EXOCYTOSIS
GO_SECRETION
GO_INTERFERON_GAMMA_MEDIATED_SIGNALING_PATHWAY
GO_RIBOSOME_ASSEMBLY
GO_RRNA_METABOLIC_PROCESS
GO_RNA_PROCESSING
GO_CELLULAR_MACROMOLECULE_LOCALIZATION
GO_NCRNA_PROCESSING
GO_NCRNA_METABOLIC_PROCESS
GO_RIBOSOMAL_SMALL_SUBUNIT_BIOGENESIS
GO_RIBOSOME_BIOGENESIS
GO_RIBONUCLEOPROTEIN_COMPLEX_SUBUNIT_ORGANIZATION
GO_INTERSPECIES_INTERACTION_BETWEEN_ORGANISMS
GO_INTRACELLULAR_TRANSPORT
GO_RIBONUCLEOPROTEIN_COMPLEX_BIOGENESIS
GO_CYTOPLASMIC_TRANSLATION
GO_INTRACELLULAR_PROTEIN_TRANSPORT
GO_ORGANONITROGEN_COMPOUND_BIOSYNTHETIC_PROCESS
GO_PROTEIN_LOCALIZATION_TO_ORGANELLE
GO_PROTEIN_TARGETING
GO_ESTABLISHMENT_OF_PROTEIN_LOCALIZATION_TO_ORGANELLE
GO_MACROMOLECULE_CATABOLIC_PROCESS
GO_PROTEIN_LOCALIZATION_TO_MEMBRANE
GO_ORGANIC_CYCLIC_COMPOUND_CATABOLIC_PROCESS
GO_CELLULAR_NITROGEN_COMPOUND_CATABOLIC_PROCESS
GO_MRNA_METABOLIC_PROCESS
GO_CELLULAR_MACROMOLECULE_CATABOLIC_PROCESS
GO_CELLULAR_AMIDE_METABOLIC_PROCESS
GO_VIRAL_GENE_EXPRESSION
GO_PEPTIDE_METABOLIC_PROCESS
GO_RNA_CATABOLIC_PROCESS
GO_TRANSLATIONAL_INITIATION
GO_NUCLEAR_TRANSCRIBED_MRNA_CATABOLIC_PROCESS_NONSENSE_MEDIATED_DECAY
GO_PROTEIN_TARGETING_TO_MEMBRANE
GO_AMIDE_BIOSYNTHETIC_PROCESS
GO_COTRANSLATIONAL_PROTEIN_TARGETING_TO_MEMBRANE
GO_ESTABLISHMENT_OF_PROTEIN_LOCALIZATION_TO_ENDOPLASMIC_RETICULUM
GO_PROTEIN_LOCALIZATION_TO_ENDOPLASMIC_RETICULUM
GO_ESTABLISHMENT_OF_PROTEIN_LOCALIZATION_TO_MEMBRANE
GO_PEPTIDE_BIOSYNTHETIC_PROCESS
GO_NUCLEAR_TRANSCRIBED_MRNA_CATABOLIC_PROCESS
−2
−1
0
1
2
Normalized enrichment score
Decreased in stress
Increased in stress

Figure S8

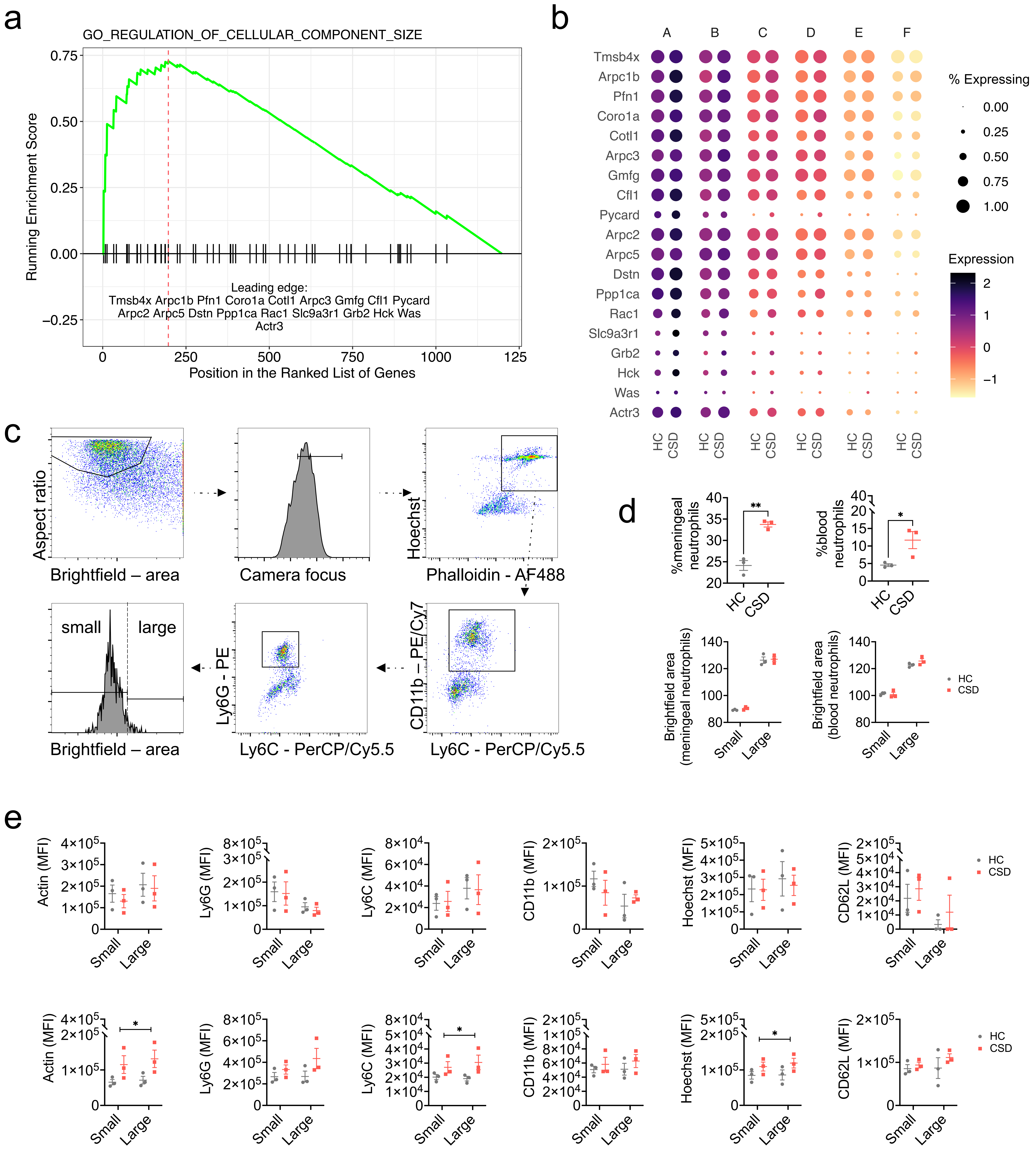
a
GO_REGULATION_OF_CELLULAR_COMPONENT_SIZE
Running Enrichment Score
Position in the Ranked List of Genes
Leading edge:
Tmsb4x Arpc1b Pfn1 Coro1a Cotl1 Arpc3 Gmfg Cfl1 Pycard
Arpc2 Arpc5 Dstn Ppp1ca Rac1 Slc9a3r1 Grb2 Hck Was
Actr3
b
A B C D E F
Tmsb4x
Arpc1b
Pfn1
Coro1a
Cotl1
Arpc3
Gmfg
Cfl1
Pycard
Arpc2
Arpc5
Dstn
Ppp1ca
Rac1
Slc9a3r1
Grb2
Hck
Was
Actr3
HC CSD
% Expressing
Expression
c
Aspect ratio
Brightfield – area
Camera focus
Hoechst
Phalloidin - AF488
small
large
Ly6G - PE
Ly6C - PerCP/Cy5.5
CD11b – PE/Cy7
d
%meningeal neutrophils
%blood neutrophils
Brightfield area (meningeal neutrophils)
Brightfield area (blood neutrophils)
Small
Large
HC
CSD
e
Actin (MFI)
Ly6G (MFI)
Ly6C (MFI)
CD11b (MFI)
Hoechst (MFI)
CD62L (MFI)

Figure S9

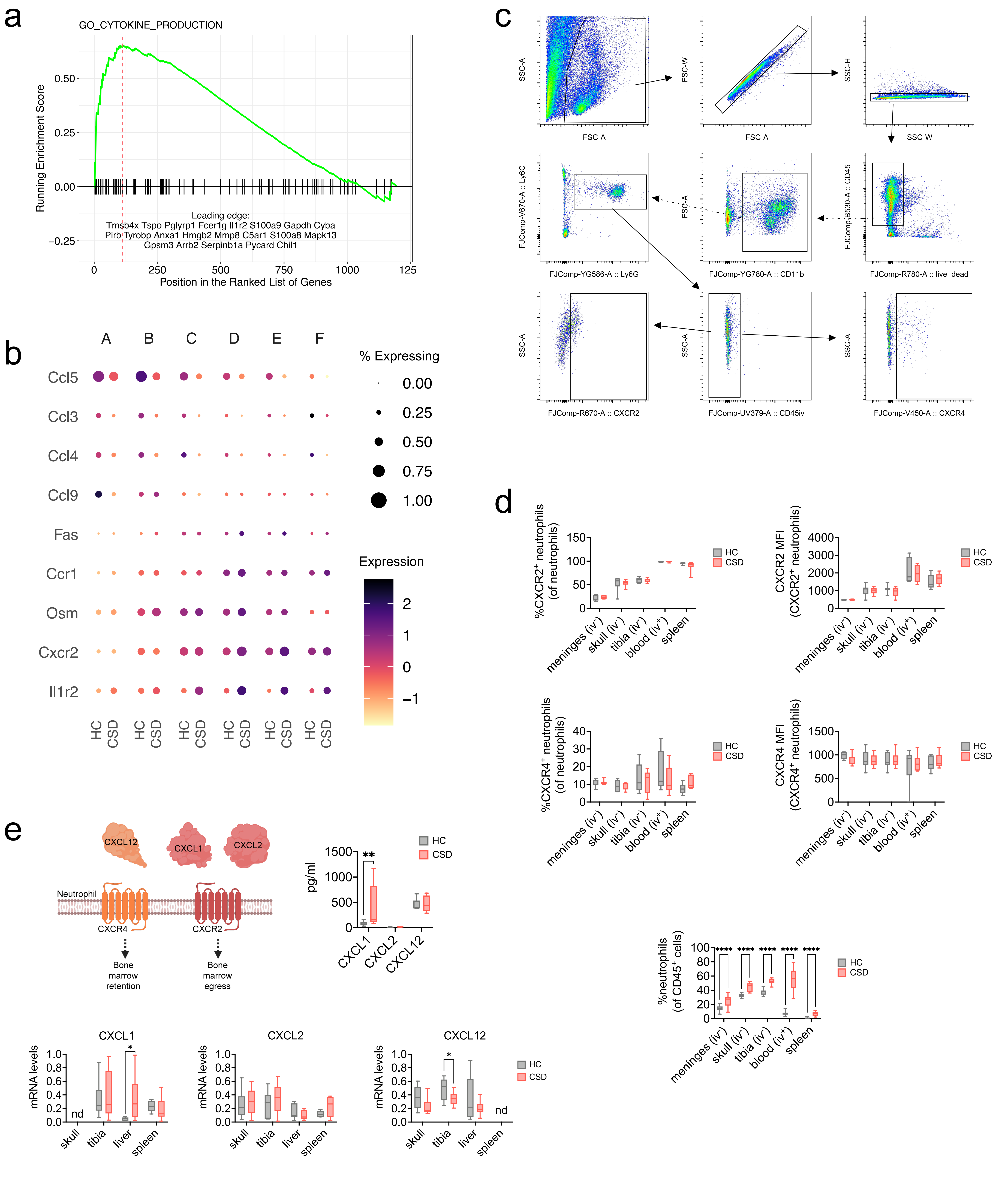

a
GO_CYTOKINE_PRODUCTION
Running Enrichment Score
Position in the Ranked List of Genes
Leading edge:
Tmsb4x Tspo Pglyrp1 Fcer1g Il1r2 S100a9 Gapdh Cyba
Pirb Tyrobp Anxa1 Hmgb2 Mmp8 C5ar1 S100a8 Mapk13
Gpsm3 Arrb2 Serpinb1a Pycard Chil1
b
A B C D E F
Ccl5
Ccl3
Ccl4
Ccl9
Fas
Ccr1
Osm
Cxcr2
Il1r2
HC CSD
% Expressing
Expression
c
SSC-A
FSC-A
FSC-W
SSC-H
SSC-W
FJComp-V670-A :: Ly6C
FJComp-YG586-A :: Ly6G
FJComp-YG780-A :: CD11b
FJComp-B530-A :: CD45
FJComp-R780-A :: live_dead
FJComp-R670-A :: CXCR2
FJComp-UV379-A :: CD45iv
FJComp-V450-A :: CXCR4
d
%CXCR2⁺ neutrophils (of neutrophils)
CXCR2 MFI (CXCR2⁺ neutrophils)
%CXCR4⁺ neutrophils (of neutrophils)
CXCR4 MFI (CXCR4⁺ neutrophils)
meninges (iv⁻)
skull (iv⁻)
tibia (iv⁻)
blood (iv⁺)
spleen
HC
CSD
e
CXCL12
CXCL1
CXCL2
Neutrophil
CXCR4
CXCR2
Bone marrow retention
Bone marrow egress
pg/ml
CXCL1
CXCL2
CXCL12
%neutrophils (of CD45⁺ cells)
****
mRNA levels
skull
tibia
liver
spleen
nd
*


Figure S10

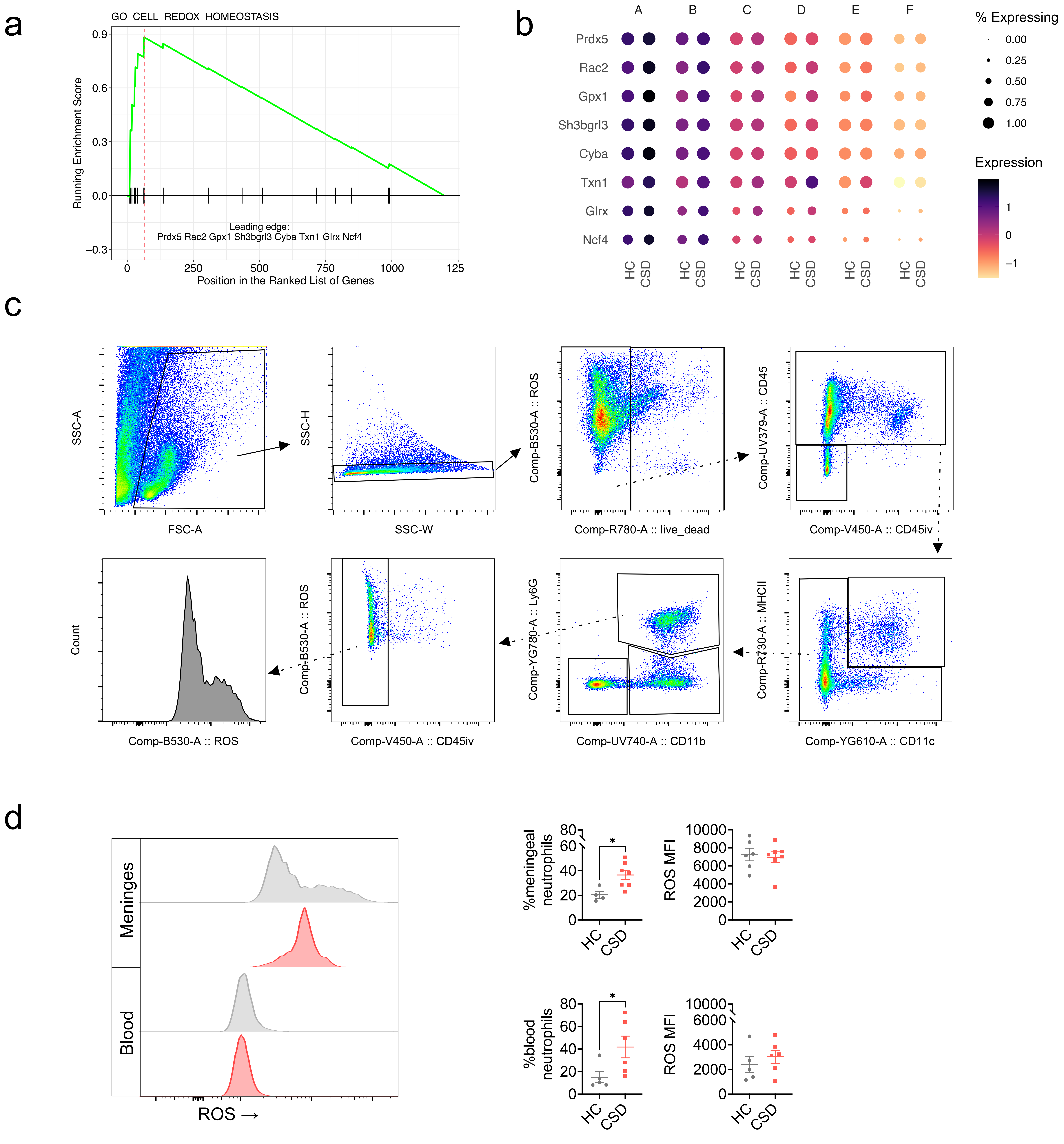
a
GO_CELL_REDOX_HOMEOSTASIS
Running Enrichment Score
Leading edge:
Prdx5 Rac2 Gpx1 Sh3bgrl3 Cyba Txn1 Glrx Ncf4
Position in the Ranked List of Genes
b
% Expressing
Expression
Prdx5
Rac2
Gpx1
Sh3bgrl3
Cyba
Txn1
Glrx
Ncf4
HC
CSD
c
SSC-A
FSC-A
SSC-H
SSC-W
Comp-B530-A :: ROS
Comp-R780-A :: live_dead
Comp-UV379-A :: CD45
Comp-V450-A :: CD45iv
Comp-R730-A :: MHCII
Comp-YG610-A :: CD11c
Comp-YG780-A :: Ly6G
Comp-UV740-A :: CD11b
Count
d
Meninges
Blood
ROS →
%meningeal neutrophils
%blood neutrophils
ROS MFI

Figure S11

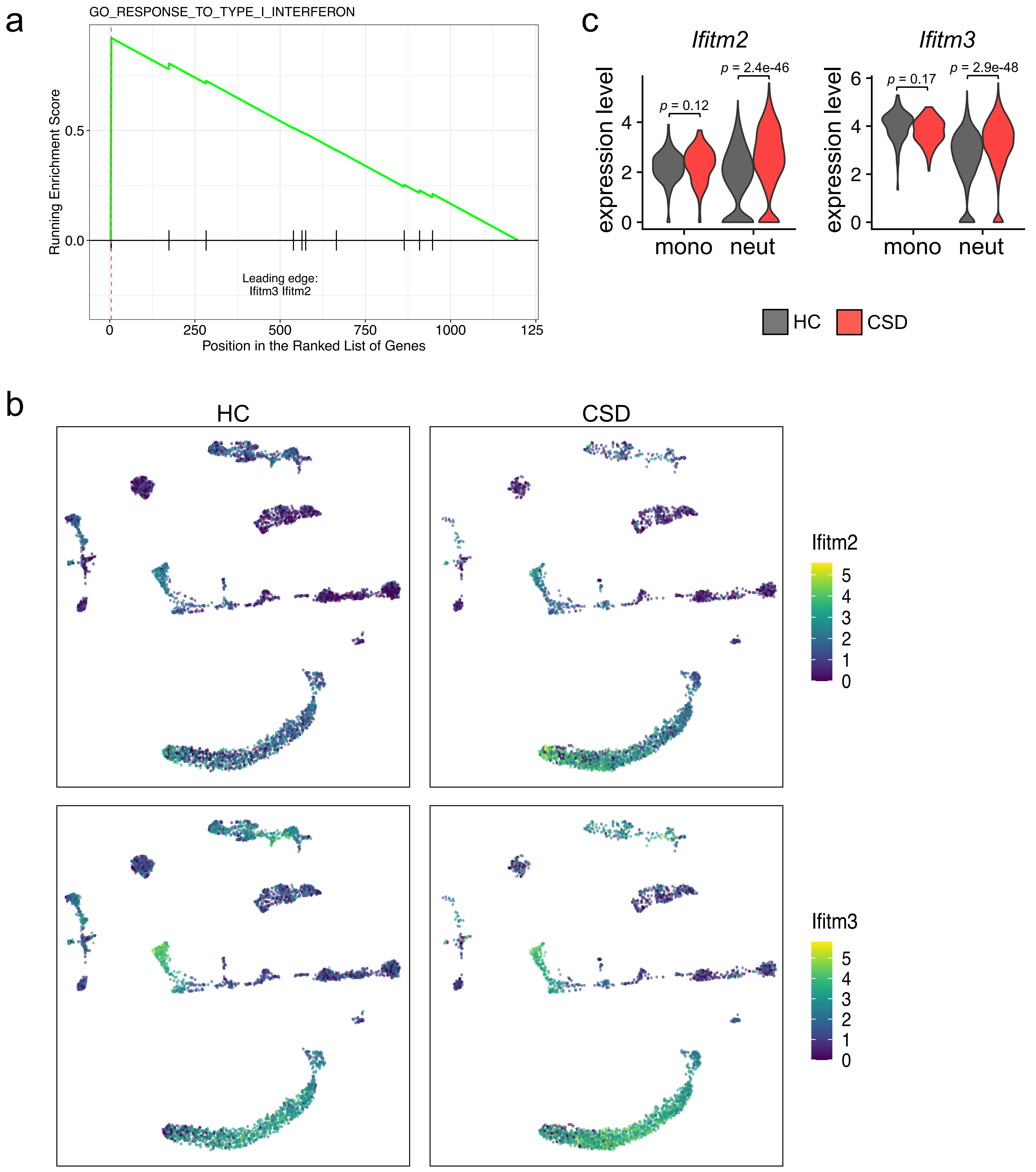
a
GO_RESPONSE_TO_TYPE_I_INTERFERON
Running Enrichment Score
0.5
0.0
Leading edge:
Ifitm3 Ifitm2
0
250
500
750
1000
125
Position in the Ranked List of Genes
c
Ifitm2
Ifitm3
expression level
p = 0.12
p = 2.4e-46
p = 0.17
p = 2.9e-48
mono
neut
HC
CSD
b
HC
CSD
Ifitm2
Ifitm3
0
1
2
3
4
5
6

Figure S12

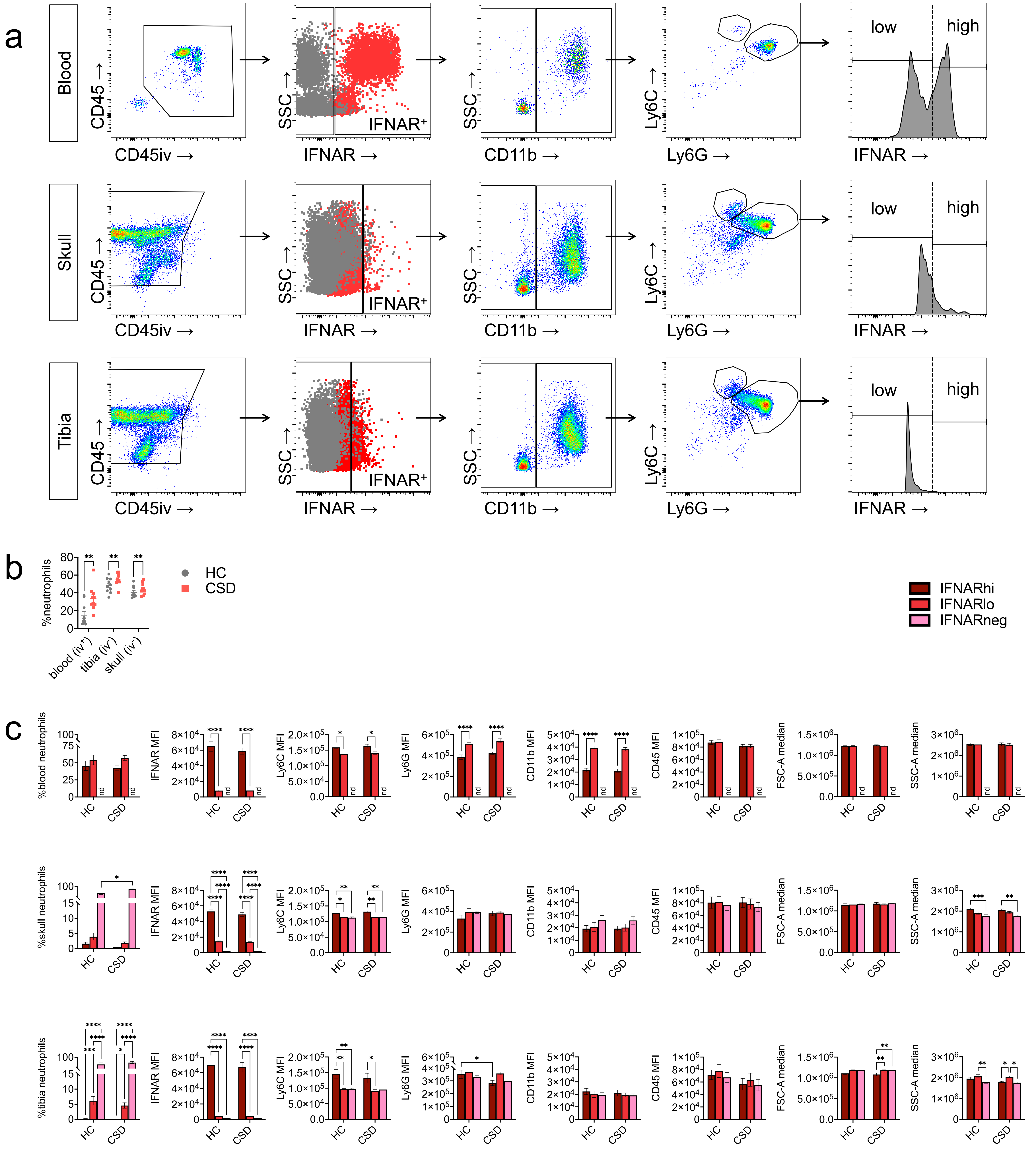
a
Blood
Skull
Tibia
CD45 →
CD45iv →
SSC →
IFNAR →
IFNAR⁺
CD11b →
Ly6C →
Ly6G →
low
high
b
%neutrophils
HC
CSD
blood (iv⁺)
tibia (iv⁻)
skull (iv⁻)
c
IFNARhi
IFNARlo
IFNARneg
%blood neutrophils
%skull neutrophils
%tibia neutrophils
IFNAR MFI
Ly6C MFI
Ly6G MFI
CD11b MFI
CD45 MFI
FSC-A median
SSC-A median
nd

Figure S13

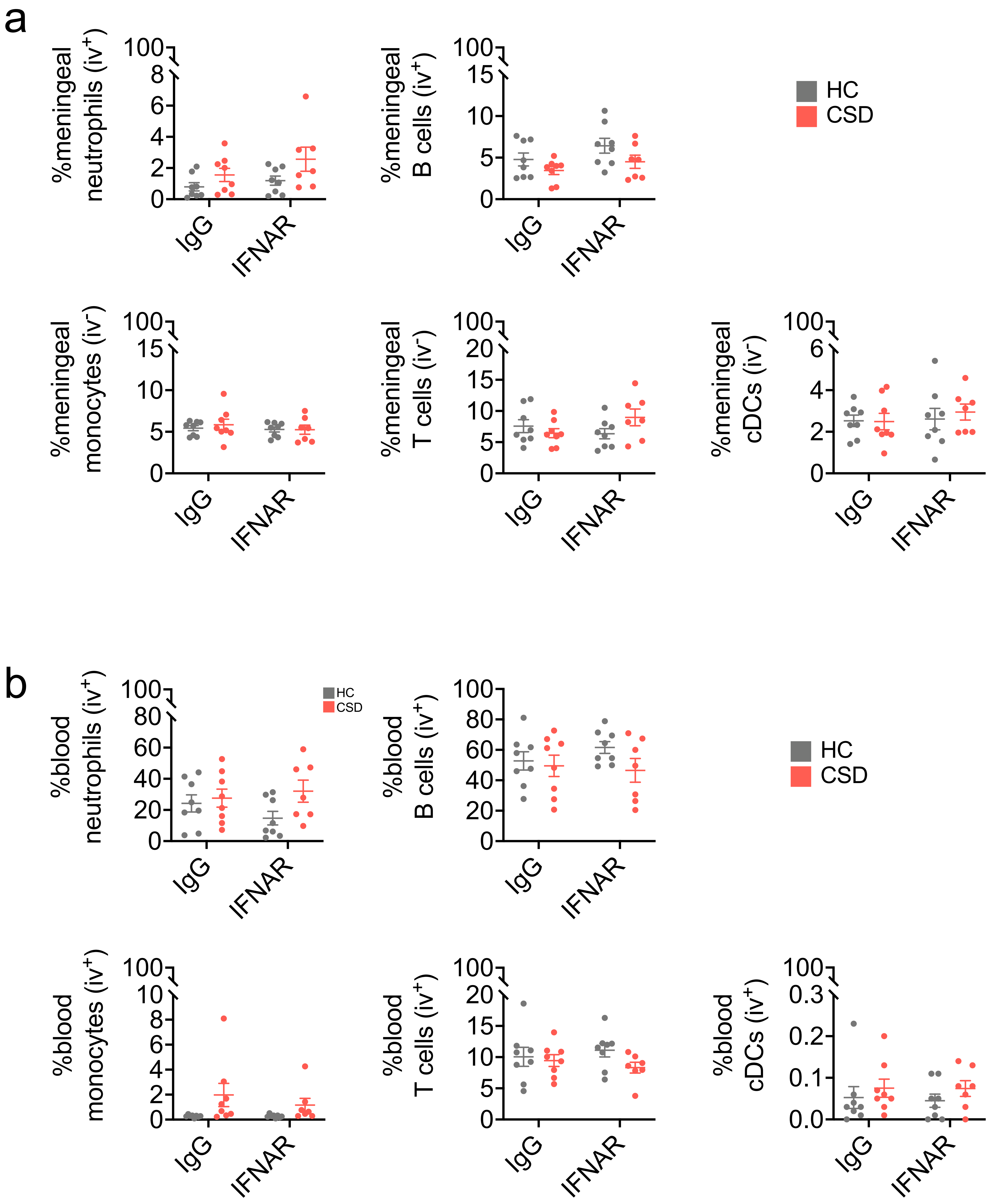
a
%meningeal neutrophils (iv+)
%meningeal B cells (iv+)
HC
CSD
%meningeal monocytes (iv-)
%meningeal T cells (iv-)
%meningeal cDCs (iv-)
IgG
IFNAR
b
%blood neutrophils (iv+)
%blood B cells (iv+)
%blood monocytes (iv+)
%blood T cells (iv+)
%blood cDCs (iv+)

Figure S14

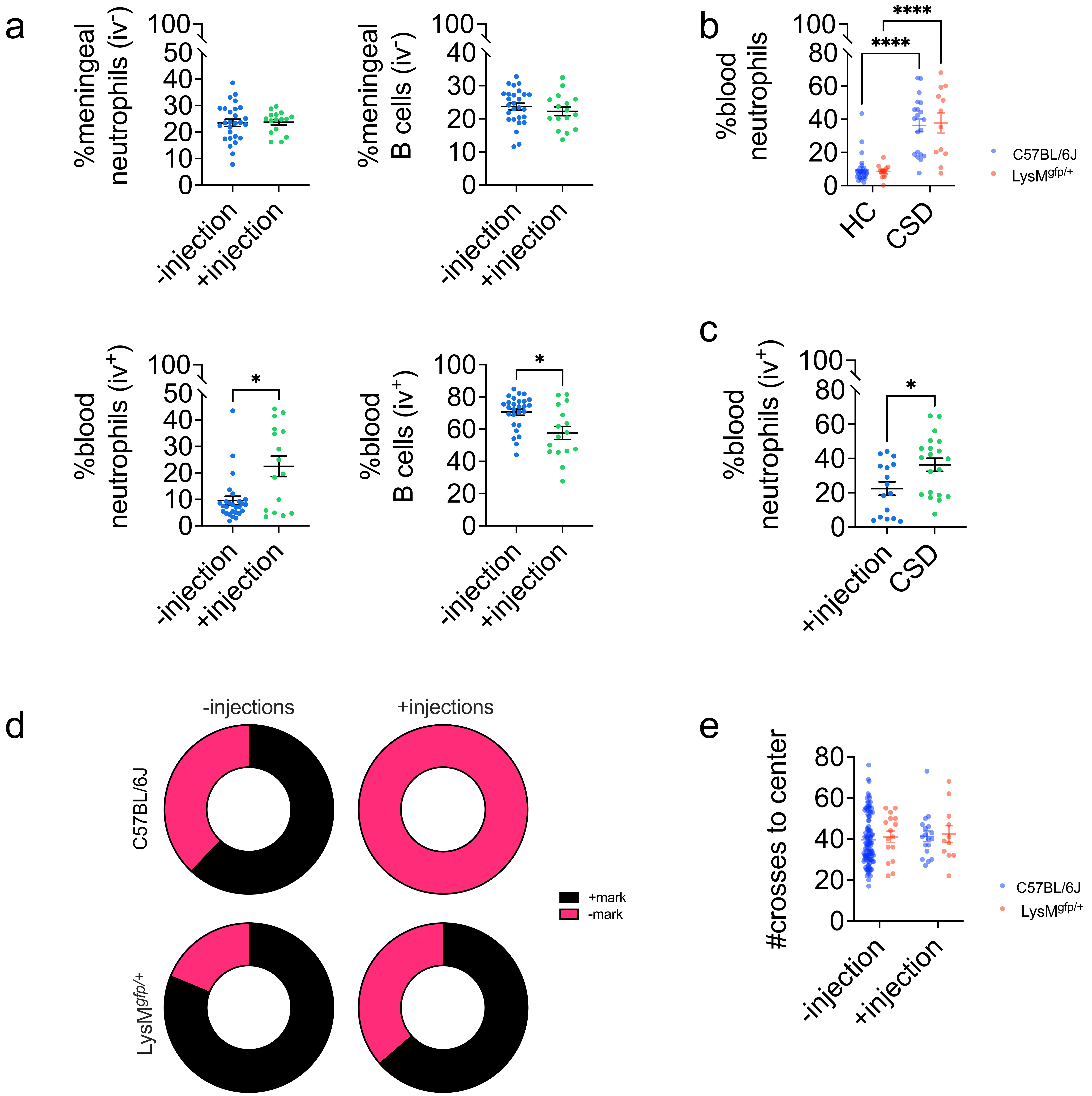
a
%meningeal neutrophils (iv$^-$)
100
50
40
30
20
10
0
-injection
+injection
%meningeal B cells (iv$^-$)
100
40
30
20
10
0
-injection
+injection
%blood neutrophils (iv$^+$)
100
50
40
30
20
10
0
*
-injection
+injection
%blood B cells (iv$^+$)
100
80
60
40
20
0
*
-injection
+injection
b
%blood neutrophils
100
80
60
40
20
0
****
****
HC
CSD
C57BL/6J
LysM$^{gfp/+}$
c
%blood neutrophils (iv$^+$)
100
80
60
40
20
0
*
+injection
CSD
d
-injections
+injections
C57BL/6J
LysM$^{gfp/+}$
+mark
-mark
e
#crosses to center
80
60
40
20
0
-injection
+injection
C57BL/6J
LysM$^{gfp/+}$

Figure S15
